# Quaternary climate change explains global patterns of tree beta-diversity

**DOI:** 10.1101/2020.11.14.382846

**Authors:** Wu-Bing Xu, Wen-Yong Guo, Josep M. Serra-Diaz, Franziska Schrodt, Wolf L. Eiserhardt, Brian J. Enquist, Brian S. Maitner, Cory Merow, Cyrille Violle, Madhur Anand, Michaël Belluau, Hans Henrik Bruun, Chaeho Byun, Jane A. Catford, Bruno E. L. Cerabolini, Eduardo Chacón-Madrigal, Daniela Ciccarelli, Johannes H. C. Cornelissen, Anh Tuan Dang-Le, Angel de Frutos, Arildo S. Dias, Aelton B. Giroldo, Alvaro G. Gutiérrez, Wesley Hattingh, Tianhua He, Peter Hietz, Nate Hough-Snee, Steven Jansen, Jens Kattge, Benjamin Komac, Nathan Kraft, Koen Kramer, Sandra Lavorel, Christopher H. Lusk, Adam R. Martin, Ke-Ping Ma, Maurizio Mencuccini, Sean T. Michaletz, Vanessa Minden, Akira S. Mori, Ülo Niinemets, Yusuke Onoda, Renske E. Onstein, Josep Peñuelas, Valério D. Pillar, Jan Pisek, Matthew J. Pound, Bjorn J.M. Robroek, Brandon Schamp, Martijn Slot, Ênio Sosinski, Nadejda A. Soudzilovskaia, Nelson Thiffault, Peter van Bodegom, Fons van der Plas, Jingming Zheng, Jens-Christian Svenning, Alejandro Ordonez

## Abstract

Both historical and contemporary environmental conditions determine present biodiversity patterns, but their relative importance is not well understood. One way to disentangle their relative effects is to assess how different dimensions of beta-diversity relate to past climatic changes, i.e., taxonomic, phylogenetic and functional compositional dissimilarity, and their components generated by replacement of species, lineages and traits (turnover) and richness changes (nestedness). Here, we quantify global patterns of each of these aspects of beta-diversity among neighboring sites for angiosperm trees using the most extensive global database of tree species-distributions (43,635 species). We found that temperature change since the Last Glacial Maximum (LGM) was the major influence on both turnover and nestedness components of beta-diversity, with a negative correlation to turnover and a positive correlation to nestedness. Moreover, phylogenetic and functional nestedness was higher than expected from taxonomic beta-diversity in regions that experienced large temperature changes since the LGM. This pattern reflects relatively greater losses of phylogenetic and functional diversity in species-poor assemblages, possibly caused by phylogenetically and functionally selective species extinction and recolonization during glacial-interglacial oscillations. Our results send a strong warning that rapid anthropogenic climate change is likely to result in a long-lasting phylogenetic and functional compositional simplification, potentially impairing forest ecosystem functioning.

## Introduction

Understanding spatial variation in biodiversity and how biodiversity responds to climate change are critical issues in ecology ^1–5^. Several studies focusing on these issues examine paleoclimatic legacies in current biodiversity patterns ^2,6–14^. These studies have shown that past climate changes have affected present species distributions and biodiversity patterns by driving extinctions, colonizations, range shifts, and diversification (as reviewed in ref. ^10^). However, these findings are primarily based on analyses of spatial patterns of local species richness (alpha-diversity), spatial variability in species composition among sites (beta-diversity) at large scales have received less attention ^10,14–19^. Because species richness is strongly determined by contemporary climatic conditions ^20,21^, which are usually correlated with paleoclimates, the extent to which historical environmental conditions shape present-day biodiversity is still under debate ^22–24^. Compared to alpha-diversity, beta-diversity considers not only the number of species but also their identities, it thus provides a way to examine how biodiversity changes across space and can enhance understanding of processes shaping biodiversity patterns ^14,17–19,25^.

To detect processes of past climate change shaping beta-diversity, it is important to partition beta-diversity into two additive components: spatial species turnover and nestedness of assemblages ^15,19,25–28^. Spatial species turnover reflects species replacement between sites, while nestedness describes the extent to which depauperate assemblages are subsets of richer ones, reflecting species loss across sites ^25^. Compared to regions with unstable climates, climatically stable regions (such as the southwest of China in Fig. 1a) have experienced lower rates of species extinction and higher rates of speciation ^6^, which leads to more species with small ranges ^29–32^. As a result, spatial species turnover is expected to be the dominant component of beta-diversity in climatically stable regions (Fig. 1b). In contrast, climatically unstable regions have experienced more local extinctions, and many extant species are recolonizers from neighboring regions after glaciations ^6^. Because postglacial recolonization lags behind the change in suitable climatic conditions ^8,33^, species richness tends to decrease from glacial refugia towards glaciated regions, leading to nestedness patterns in community compositions ^15,27,34,35^. Therefore, the nestedness component of beta-diversity is likely to be high in climatically unstable regions (Fig. 1c).

**Fig. 1.**
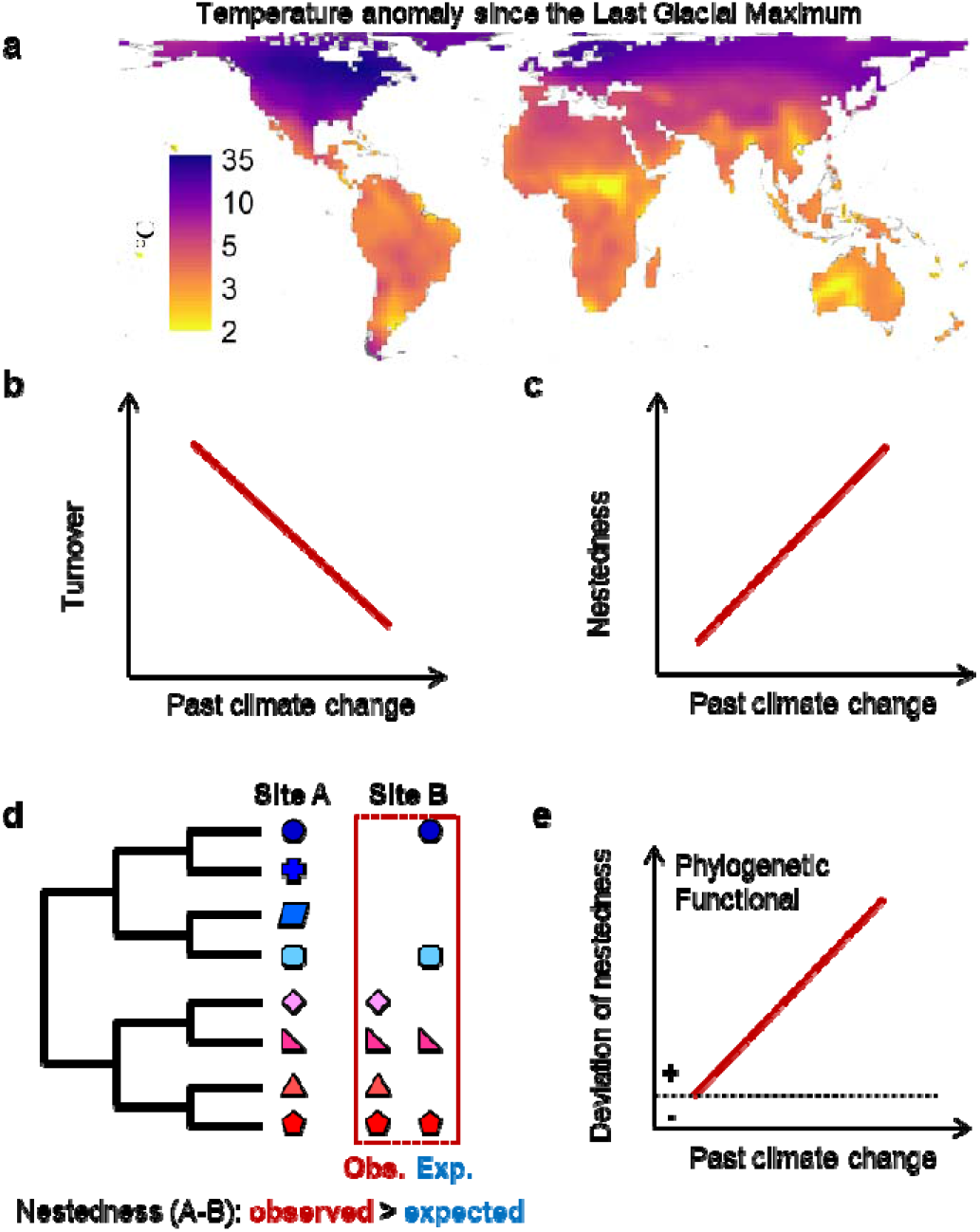
Hypothetical associations between taxonomic, phylogenetic or functional beta-diversity and past climate change. **a**, Global map of temperature anomaly since the Last Glacial Maximum (LGM). Beta-diversity was measured as intra-regional compositional dissimilarity among 3×3 grid-cells of 200 km using Sørensen multiple-site dissimilarity. It was partitioned into the turnover component as Simpson multiple-site dissimilarity, and the nestedness component as the difference between Sørensen and Simpson dissimilarities. **b**, The turnover component captures the replacement of species, lineages or traits from one site to the other. It is predicted to decrease with past climate change. **c**, The nestedness component captures the losses of species, lineages or traits between assemblages. It is predicted to increase with past climate change. **d**, Differences in the experienced past climatic change between site A and B result in relatively stable sites (A) having a higher richness than unstable areas (B). However, how differences in species losses affect phylogenetic and functional nestedness depends on the selectiveness of lineages (branches) and traits (colours) in the losses of species (geometric shapes). **e**, If glaciation-driven extinctions and post-glacial recolonization are phylogenetic and functional selective, the deviations of observed phylogenetic and functional nestedness from a random expectation are thus positive, and increase with the magnitude of past climate change.

While a taxonomic measure of beta-diversity (TBD) and its components can indicate how species composition changes among assemblages, incorporating phylogenetic and functional information into analyses of beta-diversity provides a further process-based understanding of how biodiversity patterns are shaped ^14,18,19,28,36–38^. Phylogenetic beta-diversity (PBD) measures how deep in evolutionary time species from different assemblages have been separated, and functional beta-diversity (FBD) measures the extent to which species are functionally dissimilar among assemblages ^36,38^. Similar to taxonomic beta-diversity, phylogenetic and functional beta-diversity can also be decomposed into their turnover component due to replacement of lineages and traits; and nestedness component due to losses in lineages and functions across sites ^19,39–42^. Simultaneously considering these multiple dimensions of beta-diversity and their turnover and nestedness components can provide insights into the contemporary ecological and historical evolutionary mechanisms that structure spatial variation in biodiversity ^19,39–42^.

There is much evidence that past climate change influences phylogenetic and functional diversity, reflecting differential impacts on species according to their phylogenetic positions or functional traits (see review in ref. ^10^). Glaciation-driven extinction of temperate trees has been shown to be most common among cold-intolerant species ^9,43^. Due to time-lagged migration, dispersal capacities determine the extent to which species recolonize their climatically suitable areas after glaciations ^8,44^. Because plant traits are often strongly correlated ^45,46^, species extinction and recolonization may also be related to other traits besides cold tolerance and dispersal-related traits. These nonrandom effects may cause lower functional diversity in regions that experienced strong glacial-interglacial oscillations ^11^. When cold tolerance and dispersal-related traits are phylogenetically conserved ^9,47,48^, species extinction and recolonization during glacial-interglacial oscillations would show a strong phylogenetic signal, leading to larger than random losses of phylogenetic diversity ^9^. Among communities where nestedness is the dominant beta-diversity component, differences in phylogenetic and functional diversity between species-rich and poor assemblages would be higher than those expected based on random processes when surviving and recolonizing species in depauperate sites are phylogenetically closely related or functionally similar (Fig. 1d). The phylogenetic and functional nestedness relative to the random expectation is therefore expected to be higher in climatically unstable regions (Fig. 1e).

Despite many efforts to understand spatial distributions of tree diversity, only a few recent studies have mapped global patterns of species richness, phylogenetic and functional diversity of trees ^49–51^. To better understand the drivers of spatial variation of tree diversity and determine the potential impacts of ongoing climate change, it is essential to evaluate the influence of past climate change on global present-day patterns of tree beta-diversity. Here, we combined the most extensive global database of tree species’ distributions with information about their phylogenetic relationships and functional traits to quantify intra-regional compositional dissimilarity of angiosperm trees within 3×3 grid-cells of 200 km in three biodiversity dimensions (taxonomic, phylogenetic and functional), using a multiple-site dissimilarity approach ^52^. We partitioned all three dimensions of beta-diversity into their turnover and nestedness components. We then used temperature anomaly between the present and the Last Glacial Maximum (LGM; ~21,000 yr ago) and current climatic and topographic variables to explain the spatial variation in beta-diversity. To evaluate whether past climate change non-randomly affected species driving a higher phylogenetic and functional nestedness, we use a null model to calculate the deviation of the observed from a random expectation given the observed taxonomic beta-diversity and the regional species pool. We hypothesized that temperature anomaly since the LGM has a major influence on spatial patterns of both turnover and nestedness components of beta-diversity, but that it is negatively correlated with turnover and positively with nestedness. We also hypothesized that the observed phylogenetic and functional nestedness would be higher than expected from taxonomic beta-diversity in regions strongly affected by climate change since the LGM.

## Results

### Global patterns of tree beta-diversity

We first mapped global patterns of beta-diversity for angiosperm trees as Sørensen multiple-site dissimilarity among 3×3 grid-cells, the turnover component as Simpson multiple-site dissimilarity, and the nestedness component as the difference between Sørensen and Simpson dissimilarities ^53^. Globally, total beta-diversity and its turnover and nestedness components exhibited different spatial patterns for all three evaluated biodiversity dimensions (Fig. 2 and Supplementary Fig. 1). High total beta-diversity was mainly concentrated in southern Eurasia, Africa, the Andes, and the western United States (Fig. 2a-c). The turnover component decreased towards the poles, and its spatial patterns matched those of total beta-diversity (Fig. 2d-f and Supplementary Fig. 1d-f). By comparison, spatial patterns of the nestedness component diverged from those of the turnover component, increasing towards the poles (Fig. 2g-i and Supplementary Fig. 1g-i). Overall, the turnover component contributed more on average to total beta-diversity (Fig. 2j-l). However, the proportion of total beta-diversity contributed by the nestedness component was more than half in regions that were glaciated during the LGM, especially for phylogenetic and functional beta-diversity (Fig. 2j-l). This resulted in a positive association between the nestedness proportion and latitude (Supplementary Fig. 1j-l).

**Fig. 2.**
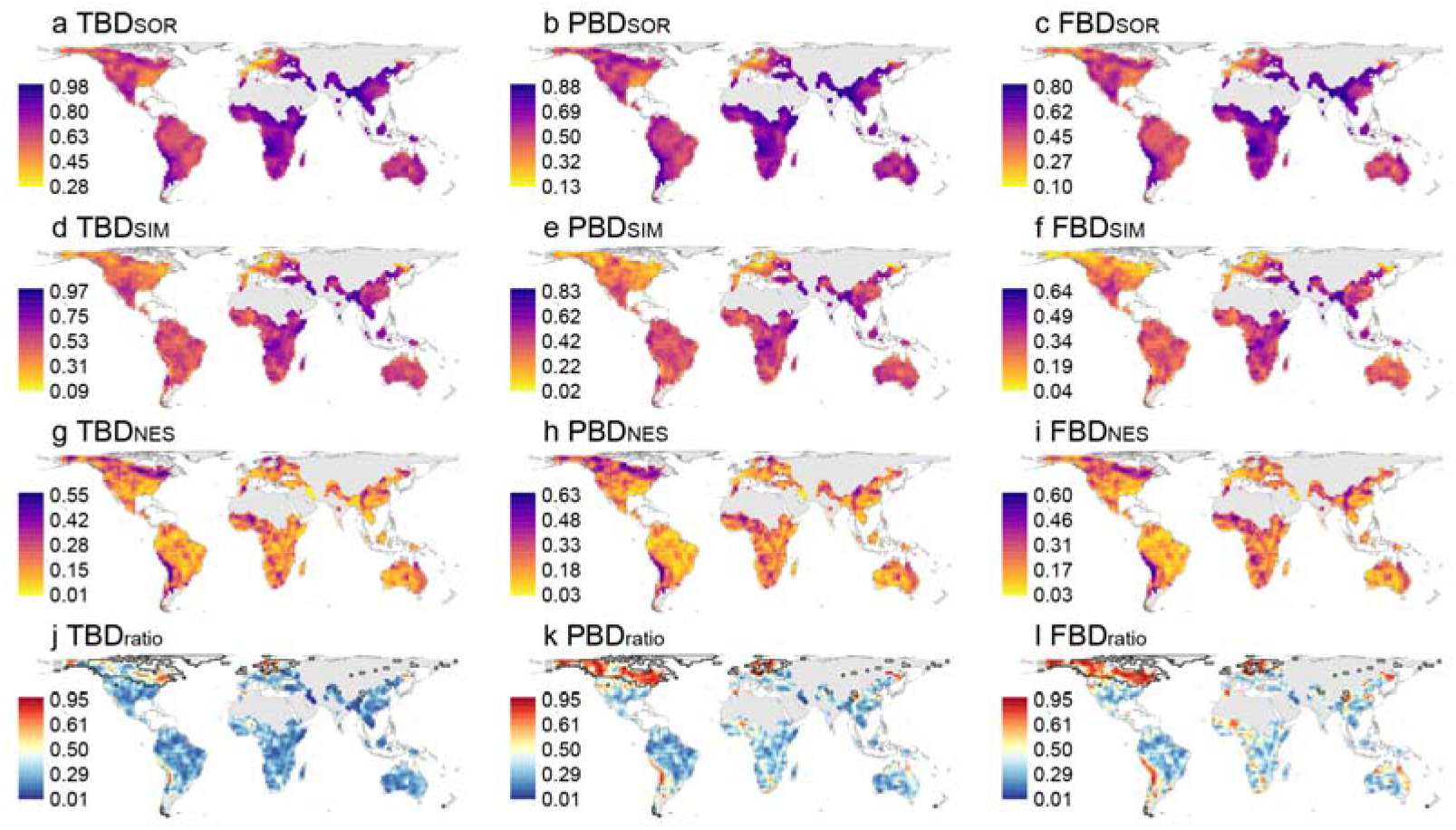
Geographic patterns of beta-diversity. **a-c**, Total beta-diversity for the taxonomic (TBD_SOR_), phylogenetic (PBD_SOR_), and functional (FBD_SOR_) dimensions; **d-f**, the component attributable to spatial turnover (TBD_SIM_; PBD_SIM_, FBD_SIM_); **g-i**, the component attributable to nestedness (TBD_NES_; PBD_NES_, FBD_NES_); **j-l**, the proportion of total beta-diversity contributed by nestedness component (TBD_ratio_; PBD_ratio_, FBD_ratio_). Black lines in j-l showed grid-cells with over half of its area covered by ice during the Last Glacial Maximum, calculated with updated Quaternary glaciation coverage maps ^95^. Grid-cells with fewer than five species or five adjacent cells are shown in gray.

### Comparison among taxonomic, phylogenetic and functional beta-diversity

Spatial patterns of total beta-diversity and its turnover and nestedness components were generally congruent among the three dimensions of biodiversity (Fig. 2), consistent with strong positive correlations between taxonomic, phylogenetic and functional beta-diversity (Supplementary Fig. 2). On average, taxonomic beta-diversity was higher than phylogenetic beta-diversity, which in turn exceeded functional beta-diversity; the same pattern was observed for the turnover component (Supplementary Fig. 2a-f). However, the nestedness component of phylogenetic and functional beta-diversity was slightly higher than that of taxonomic beta-diversity (Supplementary Fig. 2g-i), resulting in a higher relative contribution of the nestedness component for phylogenetic and functional beta-diversity than for taxonomic beta-diversity (Supplementary Fig. 2j-l).

### Associations with environmental variables

Precipitation seasonality explained most of the variation in total beta-diversity for three biodiversity dimensions (Table 1). Total taxonomic beta-diversity was also negatively associated with the temperature anomaly since the LGM, and total phylogenetic and functional beta-diversity was positively associated with mean annual temperature (Table 1). Both the turnover and nestedness components were most influenced by temperature anomaly and mean annual temperature, but they presented contrasting associations (Fig. 3; Table 1). The turnover component was negatively associated with temperature anomaly and positively with mean annual temperature for all three beta-diversity dimensions. In contrast, the nestedness component was positively associated with temperature anomaly for taxonomic and phylogenetic beta-diversity and negatively with mean annual temperature for phylogenetic beta-diversity. The relative contribution of the nestedness component showed a consistent and positive relationship with temperature anomaly and a negative association with mean annual temperature across all three dimensions of beta-diversity (Table 1).

**Fig. 3.**
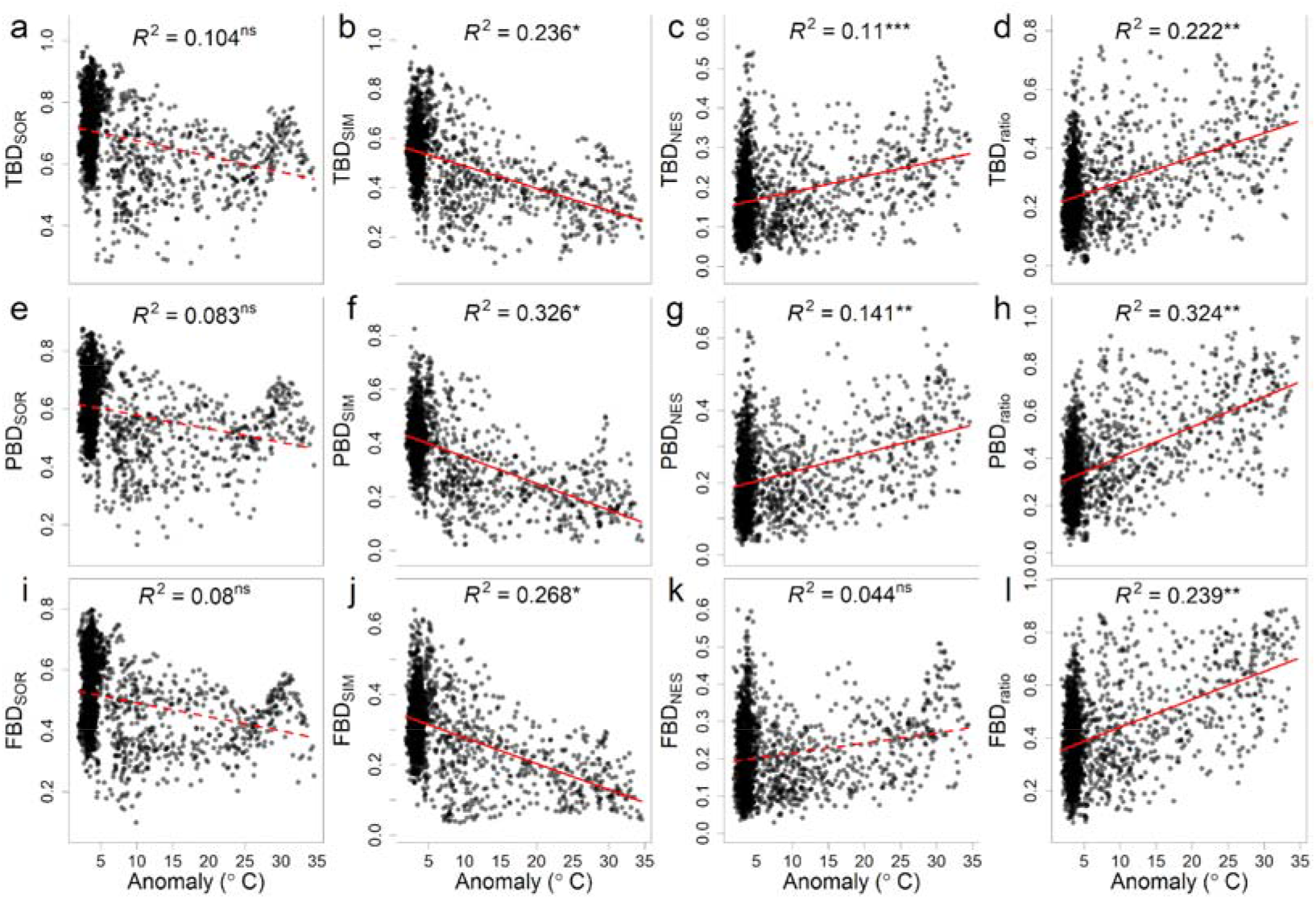
Relationships of temperature anomaly since the Last Glacial Maximum with beta-diversity. **a**,**e**,**i**, Relationships for total beta-diversity for the taxonomic (TBD_SOR_), phylogenetic (PBD_SOR_), and functional (FBD_SOR_) dimensions; **b**,**f**,**j**, relationships for the component attributable to spatial turnover (TBD_SIM_; PBD_SIM_, FBD_SIM_); **c**,**g**,**k**, relationships for the component attributable to nestedness (TBD_NES_; PBD_NES_, FBD_NES_); **d**,**h**,**l**, relationships for the proportion of total beta-diversity contributed by nestedness component (TBD_ratio_; PBD_ratio_, FBD_ratio_). The red lines were fitted with linear regressions. Significance was tested using modified *t*-test to control for spatial autocorrelation. The solid and dashed lines indicated significant and insignificant relationships, respectively. *, *P* < 0.05; **, *P* < 0.01; ***, *P* < 0.001; ns, not significant.

**Table 1.**
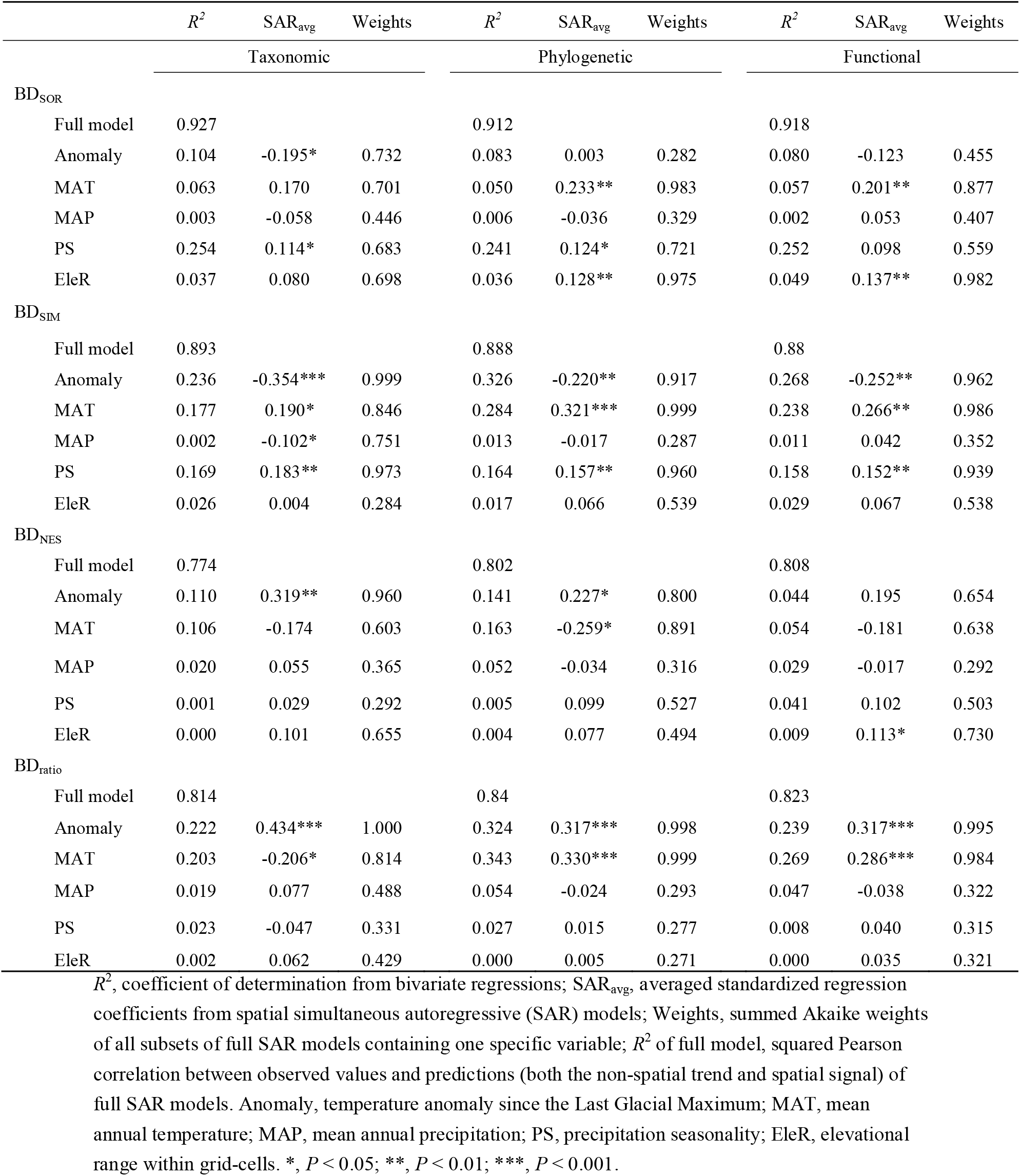
Relationships of five environmental predictors to total taxonomic, phylogenetic and functional beta-diversity (BD_SOR_) and the components attributable to spatial turnover (BD_SIM_) and nestedness (BD_NES_), and the proportion of total beta-diversity contributed by nestedness. (BD_ratio_).

| | $R^2$ | SAR<sub>avg</sub> | Weights | $R^2$ | SAR<sub>avg</sub> | Weights | $R^2$ | SAR<sub>avg</sub> | Weights |
| --- | --- | --- | --- | --- | --- | --- | --- | --- | --- |
|  | Taxonomic |  |  | Phylogenetic |  |  | Functional |  |  |
| BD<sub>SOR</sub> |  |  |  |  |  |  |  |  |  |
| Full model | 0.927 |  |  | 0.912 |  |  | 0.918 |  |  |
| Anomaly | 0.104 | -0.195* | 0.732 | 0.083 | 0.003 | 0.282 | 0.080 | -0.123 | 0.455 |
| MAT | 0.063 | 0.170 | 0.701 | 0.050 | 0.233** | 0.983 | 0.057 | 0.201** | 0.877 |
| MAP | 0.003 | -0.058 | 0.446 | 0.006 | -0.036 | 0.329 | 0.002 | 0.053 | 0.407 |
| PS | 0.254 | 0.114* | 0.683 | 0.241 | 0.124* | 0.721 | 0.252 | 0.098 | 0.559 |
| EleR | 0.037 | 0.080 | 0.698 | 0.036 | 0.128** | 0.975 | 0.049 | 0.137** | 0.982 |
| BD<sub>SIM</sub> |  |  |  |  |  |  |  |  |  |
| Full model | 0.893 |  |  | 0.888 |  |  | 0.88 |  |  |
| Anomaly | 0.236 | -0.354*** | 0.999 | 0.326 | -0.220** | 0.917 | 0.268 | -0.252** | 0.962 |
| MAT | 0.177 | 0.190* | 0.846 | 0.284 | 0.321*** | 0.999 | 0.238 | 0.266** | 0.986 |
| MAP | 0.002 | -0.102* | 0.751 | 0.013 | -0.017 | 0.287 | 0.011 | 0.042 | 0.352 |
| PS | 0.169 | 0.183** | 0.973 | 0.164 | 0.157** | 0.960 | 0.158 | 0.152** | 0.939 |
| EleR | 0.026 | 0.004 | 0.284 | 0.017 | 0.066 | 0.539 | 0.029 | 0.067 | 0.538 |
| BD<sub>NES</sub> |  |  |  |  |  |  |  |  |  |
| Full model | 0.774 |  |  | 0.802 |  |  | 0.808 |  |  |
| Anomaly | 0.110 | 0.319** | 0.960 | 0.141 | 0.227* | 0.800 | 0.044 | 0.195 | 0.654 |
| MAT | 0.106 | -0.174 | 0.603 | 0.163 | -0.259* | 0.891 | 0.054 | -0.181 | 0.638 |
| MAP | 0.020 | 0.055 | 0.365 | 0.052 | -0.034 | 0.316 | 0.029 | -0.017 | 0.292 |
| PS | 0.001 | 0.029 | 0.292 | 0.005 | 0.099 | 0.527 | 0.041 | 0.102 | 0.503 |
| EleR | 0.000 | 0.101 | 0.655 | 0.004 | 0.077 | 0.494 | 0.009 | 0.113* | 0.730 |
| BD<sub>ratio</sub> |  |  |  |  |  |  |  |  |  |
| Full model | 0.814 |  |  | 0.84 |  |  | 0.823 |  |  |
| Anomaly | 0.222 | 0.434*** | 1.000 | 0.324 | 0.317*** | 0.998 | 0.239 | 0.317*** | 0.995 |
| MAT | 0.203 | -0.206* | 0.814 | 0.343 | 0.330*** | 0.999 | 0.269 | 0.286*** | 0.984 |
| MAP | 0.019 | 0.077 | 0.488 | 0.054 | -0.024 | 0.293 | 0.047 | -0.038 | 0.322 |
| PS | 0.023 | -0.047 | 0.331 | 0.027 | 0.015 | 0.277 | 0.008 | 0.040 | 0.315 |
| EleR | 0.002 | 0.062 | 0.429 | 0.000 | 0.005 | 0.271 | 0.000 | 0.035 | 0.321 |

$R^2$ , coefficient of determination from bivariate regressions; $SAR_{avg}$ , averaged standardized regression coefficients from spatial simultaneous autoregressive (SAR) models; Weights, summed Akaike weights of all subsets of full SAR models containing one specific variable; $R^2$ of full model, squared Pearson correlation between observed values and predictions (both the non-spatial trend and spatial signal) of full SAR models. Anomaly, temperature anomaly since the Last Glacial Maximum; MAT, mean annual temperature; MAP, mean annual precipitation; PS, precipitation seasonality; EleR, elevational range within grid-cells. \*, $P < 0.05$ ; \*\*, $P < 0.01$ ; \*\*\*, $P < 0.001$ .

### The deviation of phylogenetic and functional beta-diversity

Null model analyses showed that deviations of the observed phylogenetic and functional nestedness from random expectation were not evenly distributed across space (Fig. 4a,c). North America, northern Australia, and northern and western Europe showed the largest positive deviations (yellow to red colors in Fig. 4a,c), meaning strong phylogenetically and functionally selective processes. The deviations of both phylogenetic and functional nestedness were positively correlated with temperature anomaly since the LGM (Fig. 4b,d). Temperature anomaly was the most important predictor on the spatial variation in the deviations of phylogenetic and functional nestedness (Table 2). Moreover, the deviations of phylogenetic and functional nestedness showed a strong and positive relationship with the deviations of averaged absolute differences in phylogenetic and functional diversity between all intra-regional pairwise cells (*R*^2^ = 0.622 and 0.633, respectively; Supplementary Figs. 3 and 4). This association suggests that higher-than-expected phylogenetic and functional nestedness in climatically unstable regions were attributed to higher differences in phylogenetic and functional diversity among intra-regional grid-cells due to nonrandom processes (Fig. 1c-d).

**Fig. 4.**
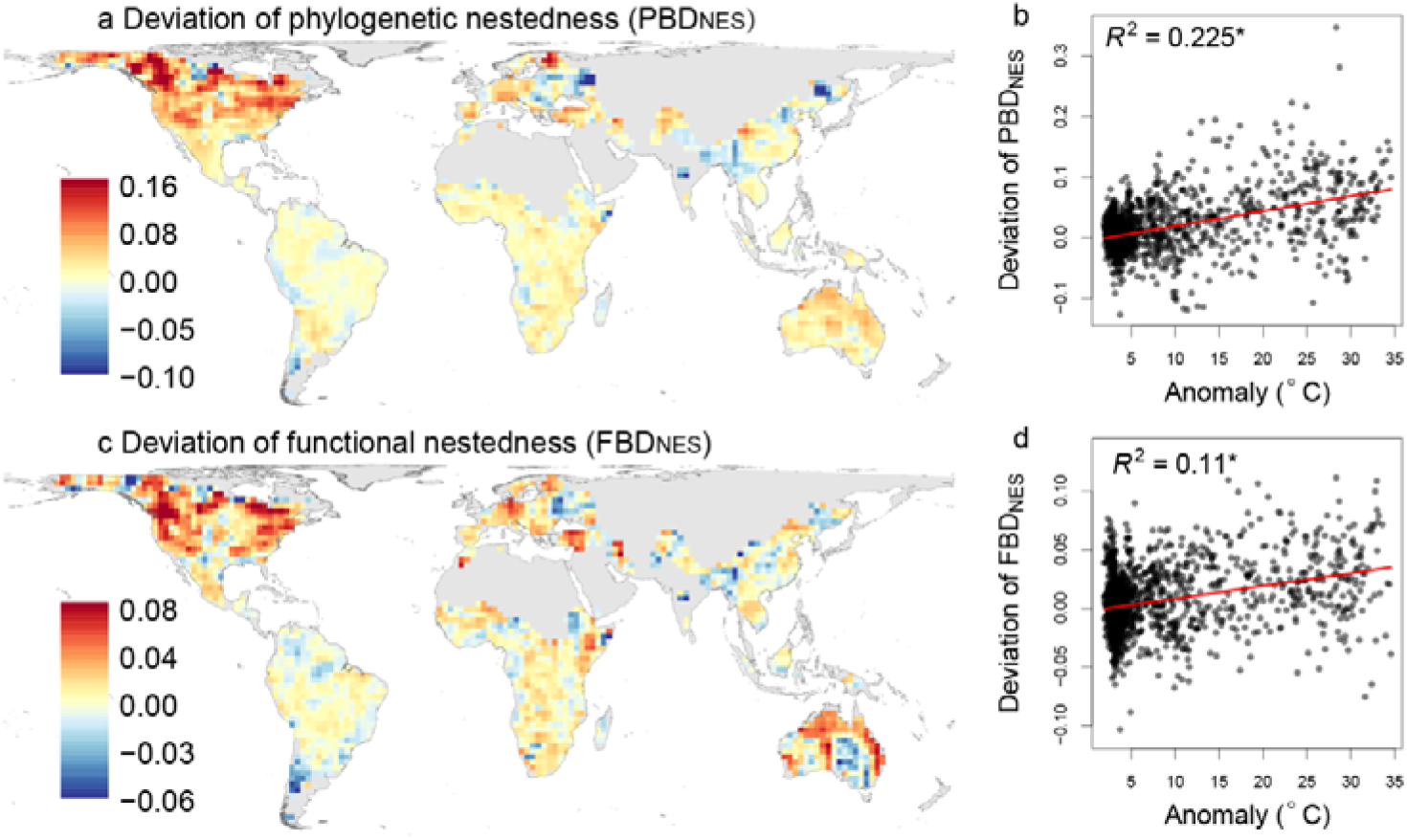
Geographic patterns of the deviation of phylogenetic and functional nestedness (a, c) and relationships with temperature anomaly since the Last Glacial Maximum (b, d). The deviation of nestedness component of phylogenetic (PBD_NES_, **a-b**) and functional (FBD_NES_, **c-d**) beta-diversity was calculated as the differences between the observed and the random expectation from taxonomic beta-diversity. The red lines in b and d were fitted with linear regressions. Significance was tested using modified *t*-test to control for spatial autocorrelation. Both relationships were significant (*P* < 0.05). Grid-cells with fewer than five species or five adjacent cells are shown in gray in a and c.

**Table 2.** Relationships of five environmental predictors with the deviations of nestedness component of phylogenetic (PBD_NES_) and functional (FBD_NES_) beta-diversity from the random expectation controlling for taxonomic beta-diversity.

| | $R^2$ | SAR<sub>avg</sub> | Weights |
| --- | --- | --- | --- |
| Deviation of PBD<sub>NES</sub> |  |  |  |
| Full model | 0.783 |  |  |
| Anomaly | 0.225 | 0.338*** | 0.999 |
| MAT | 0.166 | -0.036 | 0.297 |
| MAP | 0.049 | -0.132* | 0.886 |
| PS | 0.062 | -0.090 | 0.521 |
| EleR | 0.000 | 0.025 | 0.298 |
| Deviation of FBD<sub>NES</sub> |  |  |  |
| Full model | 0.747 |  |  |
| Anomaly | 0.110 | 0.192* | 0.799 |
| MAT | 0.050 | 0.009 | 0.332 |
| MAP | 0.020 | -0.125* | 0.754 |
| PS | 0.018 | -0.112 | 0.566 |
| EleR | 0.000 | 0.053 | 0.374 |

$R^2$ , coefficient of determination from bivariate regressions; SAR<sub>avg</sub>, averaged standardized regression coefficients from spatial simultaneous autoregressive (SAR) models; Weights, summed Akaike weights of all subsets of full SAR models containing one specific variable; $R^2$ of full model, squared Pearson correlation between observed values and predictions (both the non-spatial trend and spatial signal) of full SAR models. Anomaly, temperature anomaly since the Last Glacial Maximum; MAT, mean annual temperature; MAP, mean annual precipitation; PS, precipitation seasonality; EleR, elevational range within grid-cells. \*, $P < 0.05$ ; \*\*\*, $P < 0.001$ .

## Discussion

Using comprehensive data on angiosperm tree distributions, phylogeny, and functional traits, we showed contrasting spatial patterns of the two components of beta-diversity across three biodiversity dimensions in global tree assemblages, with the turnover component being dominant in tropical regions, and the nestedness component dominating in temperate regions. These patterns are consistent with spatial variations of beta-diversity in multiple organisms ^15,19,27,28,34,41,54^. Both patterns of the turnover and nestedness components have strong associations with temperature anomaly since the LGM, but the two relationships are in opposite directions, which reflects contrasting processes driven by past climate change on species distributions.

The high nestedness component of beta-diversity in climatically unstable regions is consistent with strong local species extinction during glaciation and incomplete postglacial recolonization from ice-age refugia ^15,27,34^. Several studies have shown that geographic accessibility from glacial refugia accounts for much of the variation in tree diversity across Europe ^13,55^. In addition, the time since deglaciation is strongly positively associated with regional species richness of vascular plants in the Arctic ^56^. These patterns are consistent with the evidence that the current distributions of many species are not in equilibrium with current climate, often due to postglacial dispersal limitation ^8,33^. Plant species with lower dispersal capacity tend to fill lower proportions of their potential ranges, and their current distributions have stronger associations with accessibility to glacial refugia ^8,57^. Such postglacial migration lags are also evident in the steeper latitudinal richness gradients of the more poorly dispersed European beetle taxa ^44^. Differential and limited dispersal capacity would result in steep rates of species losses from glacial refugia towards higher latitudes, leading to strong nestedness of community compositions in regions strongly affected by past climate change ^23^.

In contrast, the high turnover component observed in climatically stable regions could result from favored species persistence *in situ*, and the prevalence of species with small ranges ^6^. By comparison, disproportionate extinctions of small-ranged species in climatically unstable regions result in a high proportion of large-ranged species in these regions ^29–32^, decreasing the replacement of species within a given neighborhood. Therefore, many more different species would be observed among assemblages in climatically stable regions compared to those in unstable regions, resulting in higher turnover component of beta-diversity. The contrasting impacts of Quaternary climatic instability on turnover and nestedness have been reported for taxonomic beta-diversity of vertebrate groups, such as freshwater fish, amphibians, birds and mammals ^27,34,39^. By simultaneously considering taxonomic, phylogenetic and functional beta-diversity, our global analysis of trees develops previous findings and shows consistent results across three biodiversity dimensions.

Consistent with our expectation, the nestedness components of phylogenetic and functional beta-diversity are higher than the random expectation in climatically unstable regions since the LGM, reflecting nonrandom impacts of past climate change among species. This suggests that glaciation-driven local-extinction and post-glacial recolonization are most likely phylogenetically and functionally selective. Cold tolerance and dispersal capacity are thought as two important attributes affecting species survival during glaciations and recolonization to the postglacial suitable areas, respectively ^8,43,57^. These attributes are shown to have strong phylogenetic signals ^9,47,48^ and may be correlated with other functional traits as a result of biophysical constraints and trade-offs among traits ^45,46^. Within a region strongly influenced by past climate change, species-poor assemblages would have lower phylogenetic and functional diversity relative to the random expectation from the regional species pool when the extant species come from specific clades that are characterized with similar functions due to phylogenetically and functionally selective processes. The intra-regional differences in phylogenetic and functional diversity between species-rich and poor assemblages are thus expected to be higher, which is supported by our results (Supplementary Fig. 3). Because of the direct link between the intra-regional differences in phylogenetic and functional diversity and the nestedness component of beta-diversity (Supplementary Fig. 4), the higher observed differences in phylogenetic and functional diversity result in higher nestedness components of phylogenetic and functional beta-diversity in the regions that were strongly affected by Quaternary climate change.

The strong paleoclimatic legacies detected in this study imply that anthropogenic climate change is likely to have long-lasting effects on compositions of tree assemblages and thereby also on ecosystem structure and functioning ^58^. The projected geographic pattern of ongoing climate change, however, differs considerably from that during the Quaternary ^5^. The regions that are stable in paleoclimatic conditions harbouring many rare species will experience relatively faster future climate change than the past change ^5,59^. Species extinction events are projected to be more common in tropical regions due to limited dispersal and adaptation to future climate changes ^60^. Anthropogenic activities are also likely to be strong in the regions with many rare species ^59^, making climate tracking more difficult for these species. This highlights the importance of conservation tools such as assisted migration ^61^ and a well-designed network of protected areas to help species dispersal and tracking climates ^62^.

To conclude, our study shows that global patterns of tree beta-diversity in all taxonomic, phylogenetic and functional dimensions are strongly influenced by Quaternary climate change. The turnover and nestedness components of beta-diversity display distinct geographic patterns, reflecting contrasting legacies of past climate change. Further, there are higher phylogenetic and functional nestedness compared to the random expectation from taxonomic beta-diversity in regions that experienced strong past climate change, possibly caused by phylogenetically and functionally selective glaciation-driven extinction and post-glacial recolonization ^9,10^. These findings highlight the importance of climatic stability on tree diversity and suggest that anthropogenic climate change is likely to result in strong and long-lasting effects. Moreover, if species responses to anthropogenic climate changes are also phylogenetically and functionally dependent, much more phylogenetic and functional diversity will be lost than expected from the number of species going extinct, potentially leading to reduced or impaired forest ecosystem functioning ^58^.

## Methods

### Tree distributions

In this study, we used the global tree species list and distributions compiled in ref. ^63^. The tree species checklist came from the GlobalTreeSearch ^64^, which was assembled from a range of botanical publications and extended by many botanical experts. Taxonomic names were standardized using the Taxonomic Name Resolution Service (TNRS^65^), resulting in 58,100 tree species.

Tree species occurrences were compiled from five major comprehensive biodiversity infrastructures, including the Global Biodiversity Information Facility (GBIF; https://www.gbif.org), the Botanical Information and Ecological Network v.3 (BIEN^66,67^; http://bien.nceas.ucsb.edu/bien), the Latin American Seasonally Dry Tropical Forest Floristic Network (DRYFLOR^68^; http://www.dryflor.info), RAINBIO database (https://gdauby.github.io/rainbio/index.html^69^) and the Atlas of Living Australia (ALA; https://www.ala.org.au). These records were then assessed and labeled using a quality control workflow considering geographic coordinates, duplications, native ranges, geographical and environmental outliers ^63^. In this study, we used the high-quality occurrences and those records without geographic bias, which were labeled as AAA, AA, A and C in ref. ^63^. The final dataset had 46,752 species with 7,066,785 occurrences at 30 arcsec resolution.

Using the species occurrence points, we estimated species ranges based on the number of records. For species with 20 or more occurrences, ranges were estimated with alpha hulls using the R package *alphahull* ^70^. The alpha hull is a generalization of the convex hull ^71^ and allows the constructed geometric shape from a set of points to be several discrete hulls dependent on the value of the parameter alpha. For species with less than 20 occurrences or with disjunct records, a 10-km buffer was built around each point and then combined with alpha hulls. We used four alpha levels (2, 4, 6 and 10) to construct alpha hulls following recommendations in previous studies ^72,73^. Because the results based on these alpha hulls were consistent (Supplementary Figs. 5-8), our main results were based on ranges using the alpha parameter of 6 in this study. The species richness based on the range maps was strongly correlated with the prediction using forest plots and regional checklists (see ref. ^74^ for details). We noted that ranges could also be estimated by species distribution modeling using occurrences and environmental variables (e.g. ref. ^18^). In this study, we did not use the model-based ranges to avoid circular reasoning because environmental variables would be included to explain patterns of beta-diversity.

Species ranges were then rasterized to grid-cells in a resolution of 200 km with an equal-area Behrmann projection ^75^. Species assemblages in each grid-cell were defined as all species with ranges falling within the grid-cell. We chose the resolution of 200 km because biodiversity assessment at a coarse resolution can reduce under-sampling. However, we acknowledged that the patterns of beta-diversity based on these coarse tree distributions present the heterogeneity in assemblage compositions at broader spatial scales, neglecting the heterogeneity at small spatial scales. We only included angiosperms in this study, excluding gymnosperms to avoid possible divergent effects on phylogenetic and functional beta-diversity due to their striking differential evolutionary history and functional traits. We also removed grid-cells with fewer than five species to avoid potential unreliable estimates of beta-diversity. A total of 2,319 assemblages and 43,635 species were kept and used in subsequent analyses.

### Phylogenetic tree

The phylogenetic information for the tree species were extracted from the largest available phylogeny for seed plants ^76^. This comprehensive phylogeny was constructed by combining sequence data from GenBank with a backbone tree reflecting deep relationship ^77^ and adding species without molecular data but found in the Open Tree of Life (tree release 9.1 and taxonomy version 3) based on previous knowledge about phylogenetic relationships ^76^. We then reduced this phylogeny by removing any species absent in tree species list (46,752 spp.). We added some species with distributions but missing in the phylogeny using the same approach as ref. ^76^. The generated phylogeny was then pruned to contain only the angiosperm species that were used in this study.

### Functional traits

In this study, we selected eight main functional traits related to growth, survival and reproduction for functional analyses, i.e. specific leaf area, leaf area, leaf dry matter content, leaf nitrogen content, leaf phosphorus content, wood density, seed dry mass, and plant maximum height ^45^. Originally, we compiled 21 functional traits (Supplementary Table 1) from three databases, including TRY Plant Trait Database (https://www.try-db.org^78^), BIEN^66,67^ and TOPIC^79^. However, many species were missing in trait values. The trait gaps were filled by Bayesian hierarchical probabilistic matrix factorization (BHPMF), which is a robust machine learning approach imputing trait values based on the taxonomic hierarchy and correlation structure among traits ^80^. To improve estimation of missing trait values, phylogenetic information in the form of phylogenetic eigenvectors was also used as predictors ^81^. These phylogenetic eigenvectors were extracted from a principal coordinate analysis on phylogenetic distance matrix ^82^. To determine the number of phylogenetic eigenvectors included in the trait imputation, we performed preliminary analyses with increasing number of phylogenetic eigenvectors added as predictors and calculated the Root Mean Squared Error (RMSE). The RMSE was minimized as 0.087 when the first six phylogenetic eigenvectors were included in the imputation (see ref. ^74^ for details). To avoid outliers in the estimation of missing values, the maximum and minimum observed trait values were used as thresholds to constrain imputed data. For plant maximum height, a height of 2 m was used to replace imputed values lower than that considering the definition of trees in the GlobalTreeSearch database ^64^. We used all of the 21 functional traits in the imputation to maximize benefits from the correlation structure among traits.

We noted that the dataset of functional traits is limited with substantial gaps (Supplementary Table 1), which may introduce noise in the estimated functional beta-diversity. This might be the reason why lower spatial variation in functional beta-diversity was explained by environmental variables than that for phylogenetic beta-diversity in this study (Table 1). However, we believe that the functional dimension of beta-diversity can provide additional information to the phylogenetic dimension, exemplified by the detected generally lower functional turnover than phylogenetic turnover in this study.

### Beta-diversity

We used a moving window of 3×3 grid-cells of 200 km to measure intra-regional compositional heterogeneity as beta-diversity for each focal cell ^18,34^. Here, we used the multiple-site dissimilarity, rather than the average pairwise dissimilarity between focal cell and its adjacent cells because the average pairwise dissimilarity cannot reflect patterns of co-occurrence among more than two sites ^26,52^. Because differences in the number of adjacent sites could affect the multiple-site dissimilarity and some cells in the islands and along the margins of continents have few adjacent cells, we only include grid-cells having at least five adjacent occupied cells.

We calculated taxonomic, phylogenetic and functional beta-diversity using the indices from the family of Sørensen dissimilarity measures ^25,39^. Sørensen dissimilarity index is one of the most commonly used taxonomic dissimilarity indices ^83^, measuring the proportion of exclusive species among assemblages. The analogues of Sørensen dissimilarity index for phylogenetic and functional beta-diversity measure the proportion of exclusive branch lengths in a phylogenetic tree or functional dendrogram for the species among assemblages ^84^. We then partitioned these three dimensions of Sørensen dissimilarity (TBD_SOR_, PBD_SOR_ and FBD_SOR_) into two additive components due to turnover (TBD_SIM_, PBD_SIM_ and FBD_SIM_) and nestedness (TBD_NES_, PBD_NES_ and FBD_NES_) ^25,39^. The turnover component, which is the Simpson dissimilarity, reflects the effect of replacement of species or branches in a phylogeny or functional dendrogram without the impacts from differences in species richness, phylogenetic and functional diversity among sites ^25,39^. The nestedness component, which is the difference between Sørensen and Simpson dissimilarities, reflects the contribution due to differences in species richness, phylogenetic and functional diversity when species-poor assemblages are nested in species-rich assemblages ^25,39^. All calculations of beta-diversity were performed with the R package *betapart* ^53^.

Although functional beta-diversity can be calculated based on functional space ^40^, our calculation was based on a functional dendrogram, because it had similar data structure with the phylogenetic tree, making functional and phylogenetic beta-diversity comparable ^85^. To remove the correlation structure among traits, we performed principal component analysis on all eight log- and z-transformed trait values. The first three principal components (PCs) explained a high amount (83.6%) of the total variation. We thus used these three PCs to calculate the Euclidean distances among species and performed hierarchical clustering to generate the functional dendrogram.

To reflect the relative importance of components due to turnover and nestedness, we also calculated the ratio of the nestedness-resultant component relative to the total beta-diversity for all three evaluated biodiversity dimensions (TBD_ratio_, PBD_ratio_ and FBD_ratio_) ^19,27,28^. Therefore, values < 0.5 indicated that beta-diversity was determined mostly by turnover, whereas values > 0.5 suggested that the dissimilarity due to nestedness was the main component.

### Null models

As the three-dimensional beta-diversities were strongly correlated, the processes driving taxonomic beta-diversity probably also affect phylogenetic and functional beta-diversity ^28,39–41^. To investigate whether phylogenetic and functional beta-diversity were affected by processes beyond those shaping taxonomic beta-diversity, we constructed null models to calculate the random expectations of phylogenetic and functional beta-diversity given the observed taxonomic beta-diversity and regional species pool. The site-specific regional species pool was defined as all species in each focal cell and its eight adjacent cells. In the null model, the identities of species were randomly shuffled among the species pool in the phylogenetic tree and functional dendrogram. Therefore, species richness of each cell, intra-regional taxonomic beta-diversity and its turnover and nestedness components, and the pool of species, branches in the phylogenetic tree and functional dendrogram were kept constant with the observed in the null model.

Because this study focused on the processes of phylogenetically and functionally selective extinction and recolonization that could cause stronger intra-regional nestedness patterns in phylogenetic and functional compositions than expected by chance, we calculated the randomly expected phylogenetic and functional nestedness. We also calculated the observed and expected values of averaged maximum and minimum phylogenetic and functional diversity and their differences between all intra-regional pairwise cells to examine associations between phylogenetic and functional nestedness and differences in phylogenetic and functional diversity. This null model was repeated 999 times and null distributions of the phylogenetic and functional nestedness and averaged maximum and minimum phylogenetic and functional diversity and their differences were produced. We then calculated the deviations of phylogenetic and functional nestedness as the differences between the observed and the average expected values. Here, we did not use the standardized effect size because the standard deviations of expected values were strongly negatively correlated with species richness (Supplementary Fig. 9), which largely changes across the world. We also calculated the deviations of the averaged maximum and minimum phylogenetic and functional diversity and differences between pairwise cells by dividing the differences between the observed and the average expected by the observed values.

### Environmental data

To measure Quaternary climate change, we calculated the change in mean annual temperature between the present and the LGM. The mean annual temperature of the LGM was measured as the average of two estimates from GCM models CCSM4 ^86^ and MIROC-ESM ^87^ to account for variation of models. Precipitation anomaly since the LGM, however, was not used in this study because the paleoclimatic reconstruction for precipitation still remains uncertain and thus precipitation anomalies estimated from the above two GCM models differed substantially (Pearson’s *r* = 0.470).

Besides past climate change, contemporary climatic conditions and topography were also expected to shape the spatial variation of beta-diversity ^15,18,28,34,88^. We included mean annual temperature and precipitation because of their important roles affecting species distribution (Supplementary Fig. 10). Temperature seasonality and precipitation seasonality were also considered (Supplementary Fig. 10), as they were important drivers of species range size, which was directly related with beta-diversity ^31,32^. Temperature seasonality defined as the standard deviation of monthly temperature (bio4) and precipitation seasonality defined as the coefficient of variation of monthly precipitation (bio15). Topography was measured as elevational range within a cell using elevation data with a spatial resolution of 1 km^2^ (Supplementary Fig. 10).

All of the climate and elevation data used in this study came from the WorldClim 1.4 database with a resolution of 2.5 arc-min for climate ^89^. To match the scale of beta-diversity and environmental variables, environmental conditions for each cell were measured as the average of the focal cell and all of its adjacent cells ^18^.

### Statistical analyses

We used Pearson correlations to check pairwise relationships among environmental variables and also among multiple dimensional beta-diversities. To account for spatial autocorrelation in variables, a modified t-test was used to assess statistical significance ^90^. Following suggestions in ref. ^34,91^, we performed piecewise regressions to examine latitudinal patterns of beta-diversity and its turnover and nestedness components.

We then used bivariate linear regressions to investigate relationships of each metric of beta-diversity (total TBD, PBD and FBD and their turnover and nestedness components) and the deviation of phylogenetic and functional nestedness against environmental variables. We calculated the coefficient of determination (*R*^2^) to measure the importance of environmental variables. Then, multiple ordinary least squares (OLS) linear regressions were used to calculate standardized regression coefficients and determine the relative importance of environmental variables. Due to a strong correlation between mean annual temperature and temperature seasonality (Pearson’s *r* = 0.90; Supplementary Fig. 11), we excluded temperature seasonality in the multivariate analyses to avoid multicollinearity. However, residuals of all OLS models showed strong spatial autocorrelation (Supplementary Fig. 12), which could affect significance test and bias parameter estimates ^92^. To account for spatial autocorrelation, we used spatial simultaneous autoregressive (SAR) models that include a spatial weight matrix as an additional error term ^93^. A preliminary analysis was performed with a range of neighbor distances from 200 km to 1000 km in a step of 100 km and row-standardized coding style to define the spatial weight matrix. For all of our SAR models, a neighbor distance of 200 km produced a minimal Akaike information criterion value and residual spatial autocorrelation. The final SAR models successively removed spatial autocorrelation in residuals (Supplementary Fig. 12). As a supplement, the results from OLS models were provided in Supporting Information (Supplementary Table 2 and 3).

We also used Akaike-based model selection and multi-model inference to qualify the relative importance of each environmental variable by assessing all subsets of full SAR models ^94^. The Akaike weights of each model were first calculated. Then, we averaged the standardized regression coefficients for each variable across all evaluated models by weighting each value with the Akaike weight of the model that contained it. We also measured the importance of each environmental variable by summing the weights of all models including that variable.

To explore whether the regions having higher phylogenetic and functional nestedness relative to the random expectation can be attributed to their higher differences in phylogenetic and functional diversity between intra-regional pairwise cells, we investigated the bivariate linear relationships between deviations of phylogenetic and functional nestedness and deviations of differences in phylogenetic and functional diversity.

To improve linearity of regressions and normality of model residuals, mean annual precipitation and elevational range were log_10_-transformed. Although temperature anomaly was strongly right-skewed, we did not log-transform it because the residuals of models with raw anomaly closely followed normal distributions and the raw variable can explain more variation than the log-transformed variable for most metrics of beta-diversity.

All analyses were performed in R using the packages *spdep* and *spatialreg* for SAR models, *MuMIn* for multimodel inference and *SpatialPack* for modified *t*-test.

## Acknowledgement

We thank TRY contributors for sharing their data. This work was conducted as a part of the BIEN Working Group, 2008–2012. We thank all the data contributors and numerous herbaria who have contributed their data to various data compiling organizations (see the herbarium list below) for the invaluable data and support provided to BIEN. We thank the New York Botanical Garden; Missouri Botanical Garden; Utrecht Herbarium; the UNC Herbarium; and GBIF, REMIB, and SpeciesLink. The staff at CyVerse provided critical computational assistance.

We acknowledge the herbaria that contributed data to this work: A, AAH, AAS, AAU, ABH, ACAD, ACOR, AD, AFS, AK, AKPM, ALCB, ALTA, ALU, AMD, AMES, AMNH, AMO, ANGU, ANSM, ANSP, AQP, ARAN, ARIZ, AS, ASDM, ASU, AUT, AV, AWH, B, BA, BAA, BAB, BABY, BACP, BAF, BAFC, BAI, BAJ, BAL, BARC, BAS, BBB, BBS, BC, BCMEX, BCN, BCRU, BEREA, BESA, BG, BH, BHCB, BIO, BISH, BLA, BM, BOCH, BOL, BOLV, BONN, BOON, BOTU, BOUM, BPI, BR, BREM, BRI, BRIT, BRLU, BRM, BSB, BUT, C, CALI, CAN, CANB, CANU, CAS, CATA, CATIE, CAY, CBM, CDA, CDBI, CEN, CEPEC, CESJ, CGE, CGMS, CHAM, CHAPA, CHAS, CHR, CHSC, CIB, CICY, CIIDIR, CIMI, CINC, CLEMS, CLF, CMM, CMMEX, CNPO, CNS, COA, COAH, COCA, CODAGEM, COFC, COL, COLO, CONC, CORD, CP, CPAP, CPUN, CR, CRAI, CRP, CS, CSU, CSUSB, CTES, CTESN, CU, CUVC, CUZ, CVRD, DAO, DAV, DBG, DBN, DES, DLF, DNA, DPU, DR, DS, DSM, DUKE, DUSS, E, EA, EAC, EAN, EBUM, ECON, EIF, EIU, EMMA, ENCB, ER, ERA, ESA, ETH, F, FAA, FAU, FAUC, FB, FCME, FCO, FCQ, FEN, FHO, FI, FLAS, FLOR, FM, FR, FRU, FSU, FTG, FUEL, FULD, FURB, G, GAT, GB, GDA, GENT, GES, GH, GI, GLM, GMDRC, GMNHJ, GOET, GRA, GUA, GZU, H, HA, HAC, HAL, HAM, HAMAB, HAO, HAS, HASU, HB, HBG, HBR, HCIB, HEID, HGM, HIB, HIP, HNT, HO, HPL, HRCB, HRP, HSC, HSS, HU, HUA, HUAA, HUAL, HUAZ, HUCP, HUEFS, HUEM, HUFU, HUJ, HUSA, HUT, HXBH, HYO, IAA, IAC, IAN, IB, IBGE, IBK, IBSC, IBUG, ICEL, ICESI, ICN, IEA, IEB, ILL, ILLS, IMSSM, INB, INEGI, INIF, INM, INPA, IPA, IPRN, IRVC, ISC, ISKW, ISL, ISTC, ISU, IZAC, IZTA, JACA, JBAG, JBGP, JCT, JE, JEPS, JOTR, JROH, JUA, JYV, K, KIEL, KMN, KMNH, KOELN, KOR, KPM, KSC, KSTC, KSU, KTU, KU, KUN, KYO, L, LA, LAGU, LBG, LD, LE, LEB, LIL, LINC, LINN, LISE, LISI, LISU, LL, LMS, LOJA, LOMA, LP, LPAG, LPB, LPD, LPS, LSU, LSUM, LTB, LTR, LW, LYJB, LZ, M, MA, MACF, MAF, MAK, MARS, MARY, MASS, MB, MBK, MBM, MBML, MCNS, MEL, MELU, MEN, MERL, MEXU, MFA, MFU, 816 MG, MGC, MICH, MIL, MIN, MISSA, MJG, MMMN, MNHM, MNHN, MO, MOL, MOR, MPN, MPU, MPUC, MSB, MSC, MSUN, MT, MTMG, MU, MUB, MUR, MVFA, MVFQ, MVJB, MVM, MW, MY, N, NA, NAC, NAS, NCU, NE, NH, NHM, NHMC, NHT, NLH, NM, NMB, NMNL, NMR, NMSU, NSPM, NSW, NT, NU, NUM, NY, NZFRI, O, OBI, ODU, OS, OSA, OSC, OSH, OULU, OWU, OXF, P, PACA, PAMP, PAR, PASA, PDD, PE, PEL, PERTH, PEUFR, PFC, PGM, PH, PKDC, PLAT, PMA, POM, PORT, PR, PRC, PRE, PSU, PY, QCA, QCNE, QFA, QM, QRS, QUE, R, RAS, RB, RBR, REG, RELC, RFA, RIOC, RM, RNG, RSA, RYU, S, SACT, SALA, SAM, SAN, SANT, SAPS, SASK, SAV, SBBG, SBT, SCFS, SD, SDSU, SEL, SEV, SF, SFV, SGO, SI, SIU, SJRP, SJSU, SLPM, SMDB, SMF, SNM, SOM, SP, SPF, SPSF, SQF, SRFA, STL, STU, SUU, SVG, TAES, TAI, TAIF, TALL, TAM, TAMU, TAN, TASH, TEF, TENN, TEPB, TEX, TFC, TI, TKPM, TNS, TO, TOYA, TRA, TRH, TROM, TRT, TRTE, TU, TUB, U, UADY, UAM, UAMIZ, UB, UBC, UC, UCMM, UCR, UCS, UCSB, UCSC, UEC, UESC, UFG, UFMA, UFMT, UFP, UFRJ, UFRN, UFS, UGDA, UH, UI, UJAT, ULM, ULS, UME, UMO, UNA, UNB, UNCC, UNEX, UNITEC, UNL, UNM, UNR, UNSL, UPCB, UPEI, UPNA, UPS, US, USAS, USF, USJ, USM, USNC, USP, USZ, UT, UTC, UTEP, UU, UVIC, UWO, V, VAL, VALD, VDB, VEN, VIT, VMSL, VT, W, WAG, WAT, WELT, WFU, WII, WIN, WIS, WMNH, WOLL, WS, WTU, WU, XAL, YAMA, Z, ZMT, ZSS, and ZT.

WBX and AO acknowledge support from the Aarhus University Research Foundation (AUFF) Starting Grant (AUFF-F-201 8-7-8). JCS considers this work a contribution to his VILLUM Investigator project “Biodiversity Dynamics in a Changing World” funded by VILLUM FONDEN (grant 16549) and his Independent Research Fund Denmark | Natural Sciences project TREECHANGE (grant 6108-00078B), with the latter funding the development of the TREECHANGE database. JP research was supported by a Synergy grant from the European Research Council (No. ERC-2013-SyG-610028 IMBALANCE-P).

## Author contributions

WBX and AO conceived the study; WYG and JMSD and all others collated the data; WBX performed the analyses and interpreted the results; WBX and AO wrote the first draft of the manuscript; All authors contributed to interpreting results, the writing and approved the final manuscript.

