## Supplementary tables and figures for "Quaternary climate change explains global patterns of tree beta-diversity"

**Supplementary Table 1 |** The 21 functional traits compiled that were used to fill data gaps for species missing trait values based on the correlation structure among traits, taxonomic hierarchy and phylogenetic information [74]. A total of 54,020 species were included. After imputation, eight traits were selected to calculate functional beta-diversity.

| Trait name | Description | Missing species | Missing rate (%) | Selected |
|---|---|---|---|---|
| Leaf nitrogen content | Leaf nitrogen (N) content per leaf dry mass | 50,441 | 93.37 | √ |
| Wood Density | Stem specific density (SSD) | 47,608 | 88.13 | √ |
| Leaf K content | Leaf potassium (K) content per leaf dry mass | 52,778 | 97.7 | |
| Leaf P content | Leaf phosphorus (P) content per leaf dry mass | 51,652 | 95.62 | √ |
| LDMC | Leaf dry matter content | 53,227 | 98.53 | √ |
| Vegetative height | Plant maximum height | 49,521 | 91.67 | √ |
| Leaf N:P | Leaf nitrogen/phosphorus (N/P) ratio | 52,566 | 97.31 | |
| Seed dry mass | Seed dry mass | 49,348 | 91.35 | √ |
| SLA | specific leaf area (Leaf area per leaf dry mass, 1/LMA) | 52,055 | 96.36 | √ |
| LA | Leaf area (in case of compound leaves: leaflet, petiole and rachis excluded) | 53,221 | 98.52 | √ |
| SLAFM | Leaf area per leaf fresh mass (SLA based on leaf fresh mass) | 53,739 | 99.48 | |
| $LA_{compundleaf}$ | Leaf area (in case of compound leaves: leaf, petiole included) | 53,355 | 98.77 | |
| Leaf respiration rate | Leaf respiration rate per leaf area | 53,802 | 99.6 | |
| Wood N | Wood nitrogen (N) content per wood dry mass | 53,989 | 99.94 | |
| Leaf photosynthesis rate | Leaf photosynthesis rate per leaf area | 53,252 | 98.58 | |
| Seed germination rate | Seed germination rate (germination efficiency) | 53,339 | 98.74 | |
| LWR | Leaf dry mass per plant dry mass (leaf weight ratio) | 53,828 | 99.64 | |
| Stomata conductance | Stomata conductance per leaf area | 53,542 | 99.12 | |
| Fine root DM | Fine root dry mass to leaf dry mass ratio | 53,948 | 99.87 | |
| Leaf WUE | Leaf photosynthetic water use efficiency | 53,970 | 99.91 | |
| Wood DM | Wood dry mass per plant | 54,012 | 99.99 | |

**Supplementary Table 2 |** Summary results of OLS models showing relationships of five environmental predictors to total taxonomic, phylogenetic and functional beta-diversity ($BD_{SOR}$) and the components attributable to spatial turnover ($BD_{SIM}$) and nestedness ($BD_{NES}$), and the proportion of total beta-diversity contributed by nestedness ($BD_{ratio}$).

| | $R^2$ | $OLS_{avg}$ | Weights | $R^2$ | $OLS_{avg}$ | Weights | $R^2$ | $OLS_{avg}$ | Weights |
|---|---|---|---|---|---|---|---|---|---|
| | Taxonomic | | | Phylogenetic | | | Functional | | |
| $BD_{SOR}$ | | | | | | | | | |
| Full model | 0.285 | | | 0.268 | | | 0.288 | | |
| Anomaly | 0.104 | -0.088* | 0.848 | 0.083 | -0.034 | 0.363 | 0.080 | 0.041 | 0.434 |
| MAT | 0.063 | 0.089 | 0.775 | 0.050 | 0.119*** | 0.967 | 0.057 | 0.135*** | 1.000 |
| MAP | 0.003 | -0.040 | 0.592 | 0.006 | -0.065** | 0.935 | 0.002 | -0.033 | 0.500 |
| PS | 0.254 | 0.429*** | 1.000 | 0.241 | 0.416*** | 1.000 | 0.252 | 0.433*** | 1.000 |
| EleR | 0.037 | 0.133*** | 1.000 | 0.036 | 0.154*** | 1.000 | 0.049 | 0.193*** | 1.000 |
| $BD_{SIM}$ | | | | | | | | | |
| Full model | 0.309 | | | 0.395 | | | 0.352 | | |
| Anomaly | 0.236 | 0.214*** | 1.000 | 0.326 | 0.250*** | 1.000 | 0.268 | 0.164*** | 1.000 |
| MAT | 0.177 | 0.231*** | 1.000 | 0.284 | 0.326*** | 1.000 | 0.238 | 0.347*** | 1.000 |
| MAP | 0.002 | -0.058* | 0.898 | 0.013 | -0.045* | 0.769 | 0.011 | -0.041 | 0.665 |
| PS | 0.169 | 0.217*** | 1.000 | 0.164 | 0.165*** | 1.000 | 0.158 | 0.178*** | 1.000 |
| EleR | 0.026 | 0.131*** | 1.000 | 0.017 | 0.117*** | 1.000 | 0.029 | 0.177*** | 1.000 |
| $BD_{NES}$ | | | | | | | | | |
| Full model | 0.161 | | | 0.247 | | | 0.169 | | |
| Anomaly | 0.110 | 0.254*** | 1.000 | 0.141 | 0.264*** | 1.000 | 0.044 | 0.229*** | 1.000 |
| MAT | 0.106 | 0.225*** | 1.000 | 0.163 | 0.313*** | 1.000 | 0.054 | 0.187*** | 1.000 |
| MAP | 0.020 | 0.033 | 0.465 | 0.052 | -0.014 | 0.305 | 0.029 | 0.000 | 0.272 |
| PS | 0.001 | 0.211*** | 1.000 | 0.005 | 0.287*** | 1.000 | 0.041 | 0.355*** | 1.000 |
| EleR | 0.000 | -0.021 | 0.340 | 0.004 | 0.023 | 0.374 | 0.009 | 0.062* | 0.888 |
| $BD_{ratio}$ | | | | | | | | | |
| Full model | 0.253 | | | 0.397 | | | 0.318 | | |
| Anomaly | 0.222 | 0.292*** | 1.000 | 0.324 | 0.328*** | 1.000 | 0.239 | 0.260*** | 1.000 |
| MAT | 0.203 | 0.290*** | 1.000 | 0.343 | 0.391*** | 1.000 | 0.269 | 0.398*** | 1.000 |
| MAP | 0.019 | 0.049* | 0.747 | 0.054 | 0.007 | 0.281 | 0.047 | 0.021 | 0.362 |
| PS | 0.023 | 0.080** | 0.990 | 0.027 | 0.113*** | 1.000 | 0.008 | 0.165*** | 1.000 |
| EleR | 0.002 | -0.050* | 0.760 | 0.000 | -0.026 | 0.436 | 0.000 | -0.044 | 0.710 |

$R^2$, coefficient of determination from bivariate regressions; $OLS_{avg}$, averaged standardized regression coefficients from ordinary least squares (OLS) regression models; Weights, summed Akaike weights of all subsets of full OLS models containing one specific variable; $R^2$ of full model, squared Pearson correlation between observed and predicted values of full models. Anomaly, temperature anomaly since the Last Glacial Maximum; MAT, mean annual temperature; MAP, mean annual precipitation; PS, precipitation seasonality; EleR, elevational range within grid-cells. *, $P < 0.05$; **, $P < 0.01$; ***, $P < 0.001$.

**Supplementary Table 3 |** Summary results of OLS models showing relationships of five environmental predictors to the deviation of nestedness component of phylogenetic ($PBD_{NES}$) and functional ($FBD_{NES}$) beta-diversity from the random expectation controlling for taxonomic beta-diversity.

| | $R^2$ | $OLS_{avg}$ | Weights |
|---|---|---|---|
| Deviation of $PBD_{NES}$ | | | |
| Full model | 0.259 | | |
| Anomaly | 0.225 | 0.433*** | 1.000 |
| MAT | 0.166 | 0.054 | 0.498 |
| MAP | 0.049 | 0.165*** | 1.000 |
| PS | 0.062 | 0.117*** | 1.000 |
| EleR | 0.000 | 0.083*** | 0.997 |
| Devation of $FBD_{NES}$ | | | |
| Full model | 0.132 | | |
| Anomaly | 0.110 | 0.461*** | 1.000 |
| MAT | 0.050 | 0.220*** | 1.000 |
| MAP | 0.020 | 0.152*** | 1.000 |
| PS | 0.018 | -0.068** | 0.933 |
| EleR | 0.000 | 0.120*** | 1.000 |

$R^2$, coefficient of determination from bivariate regressions; $OLS_{avg}$, averaged standardized regression coefficients from ordinary least squares (OLS) regression models; Weights, summed Akaike weights of all subsets of full OLS models containing one specific variable; $R^2$ of full model, squared Pearson correlation between observed and predicted values of full models. Anomaly, temperature anomaly since the Last Glacial Maximum; MAT, mean annual temperature; MAP, mean annual precipitation; PS, precipitation seasonality; EleR, elevational range within grid-cells. ***, $P < 0.001$.

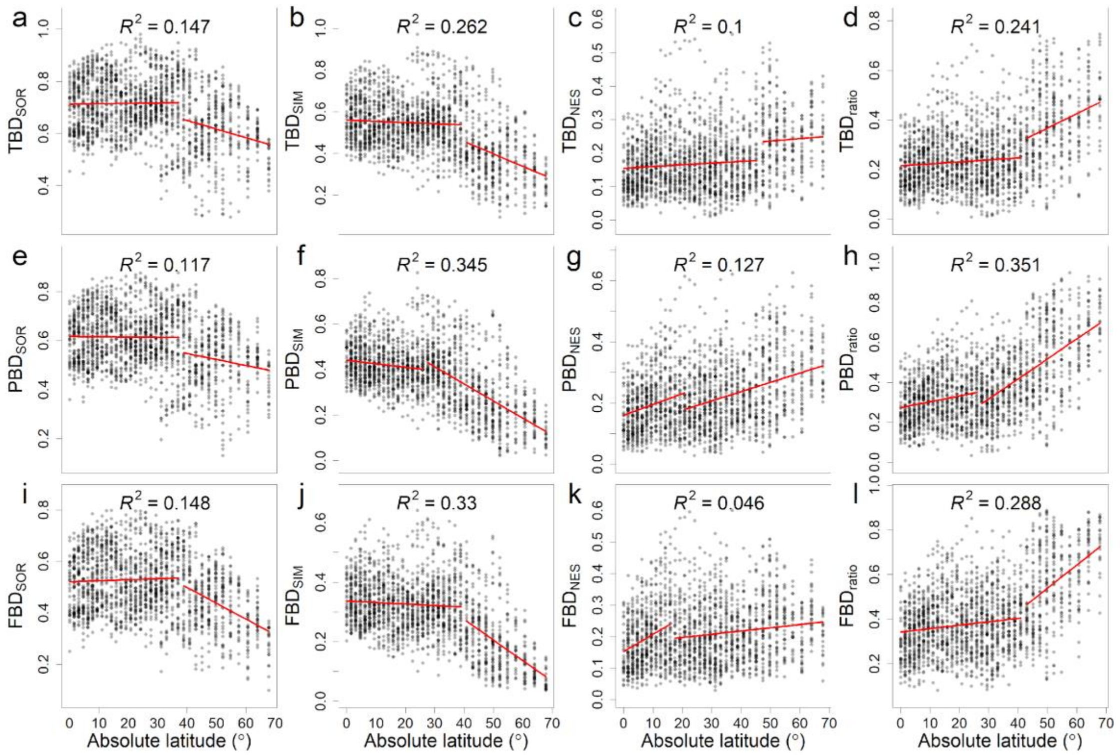


**Supplementary Fig. 1 | Latitudinal patterns of beta-diversity.** **a**,**e**,**i**, Total beta-diversity for the taxonomic ($\text{TBD}_{\text{SOR}}$), phylogenetic ($\text{PBD}_{\text{SOR}}$) and functional ($\text{FBD}_{\text{SOR}}$) dimensions; **b**,**f**,**j**, the component attributable to spatial turnover ($\text{TBD}_{\text{SIM}}$; $\text{PBD}_{\text{SIM}}$, $\text{FBD}_{\text{SIM}}$); **c**,**g**,**k** , the component attributable to nestedness ($\text{TBD}_{\text{NES}}$; $\text{PBD}_{\text{NES}}$, $\text{FBD}_{\text{NES}}$); **d**,**h**,**l**, the proportion of total beta-diversity contributed by nestedness component ($\text{TBD}_{\text{ratio}}$; $\text{PBD}_{\text{ratio}}$, $\text{FBD}_{\text{ratio}}$). The red lines were fitted with piecewise regressions.

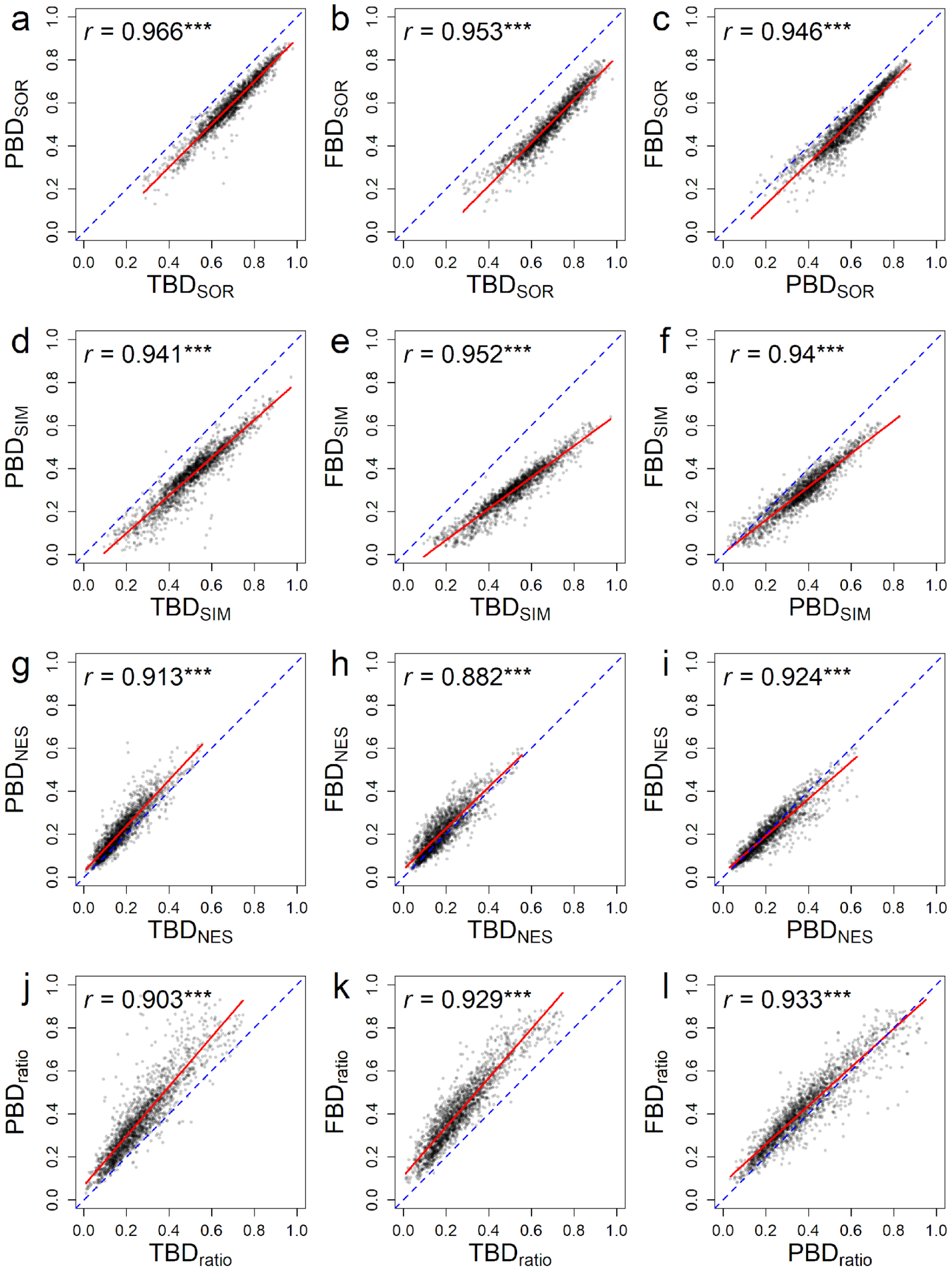


**Supplementary Fig. 2 | Relationships among taxonomic, phylogenetic and functional beta-diversity. a-c**, Relationships for the total beta-diversity for the taxonomic (TBD$_{SOR}$), phylogenetic (PBD$_{SOR}$) and functional (FBD$_{SOR}$) dimensions; **d-f**, relationships for the component attributable to spatial turnover (TBD$_{SIM}$; PBD$_{SIM}$, FBD$_{SIM}$); **g-i**, relationships for the component attributable to nestedness (TBD$_{NES}$; PBD$_{NES}$, FBD$_{NES}$;); **j-l**, relationships for the proportion of total beta-diversity contributed by nestedness component (TBD$_{ratio}$; PBD$_{ratio}$, FBD$_{ratio}$;). The red lines were fitted with linear regressions. The blue dashed lines are the identity lines. Pearson's correlation coefficients were provided. Significance was tested using modified *t*-test to control for spatial autocorrelation. ***, $P < 0.001$.

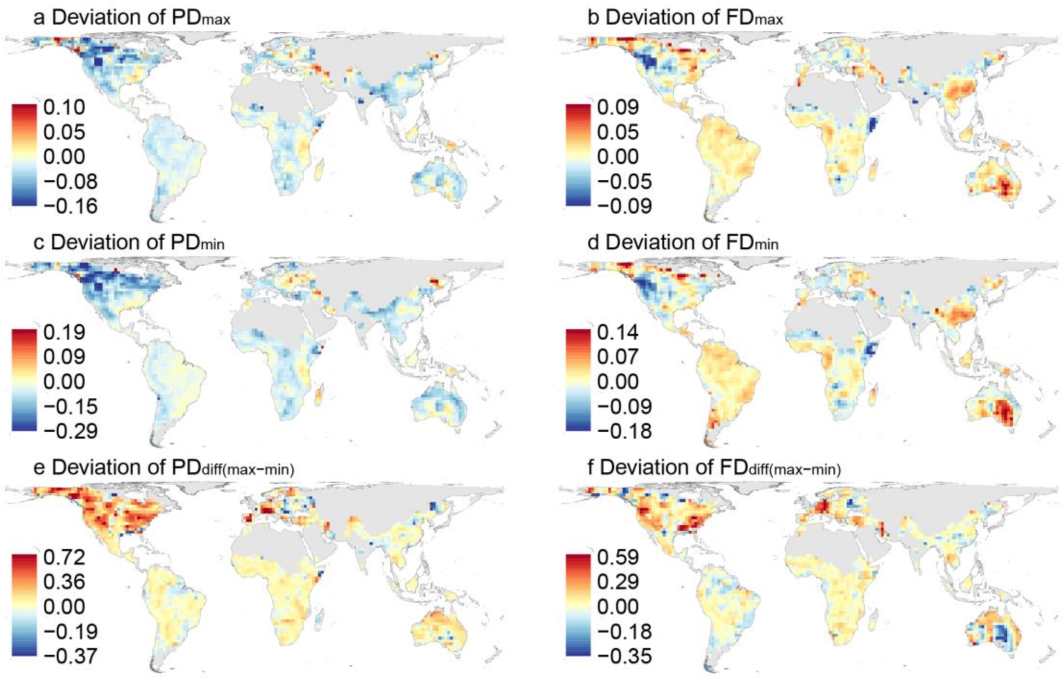


**Supplementary Fig. 3 | Geographic patterns of the deviation of intra-regional maximum, minimum, and differences in phylogenetic and functional diversity.** The intra-regional maximum (**a**, **b**) and minimum (**c**, **d**) phylogenetic ($PD_{max}$ and $PD_{min}$) and functional ($FD_{max}$ and $FD_{min}$) diversity, and their differences ($PD_{diff(max-min)}$, $FD_{diff(max-min)}$; **e**, **f**) were calculated as the averaged values of all pairwise cells in regions of 3×3 cells. All deviations were calculated as the differences between the observed and the random expectation by shuffling regional species in the phylogenetic tree and functional dendrogram. Grid-cells with fewer than five species or five adjacent cells are shown in gray.

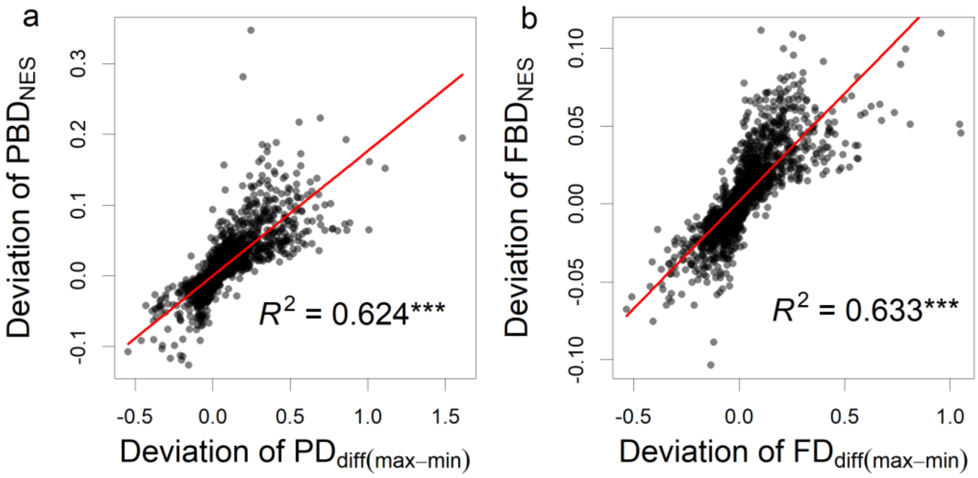


**Supplementary Fig. 4 | Relationships between deviations of phylogenetic (a) and functional (b) nestedness and deviations of intra-regional differences in phylogenetic (a)and functional (b) diversity.** The intra-regional differences in phylogenetic and functional diversity were calculated as the averaged differences between maximum and minimum phylogenetic ($PD_{diff(max-min)}$) and functional ($FD_{diff(max-min)}$) diversity of all pairwise cells in regions of 3×3 cells. All deviations were calculated as the differences between the observed and the random expectation by shuffling regional species in the phylogenetic tree and functional dendrogram. The red lines were fitted with linear regressions. Significance was tested using modified *t*-test to control for spatial autocorrelation. Both relationships were significant ($P < 0.001$).

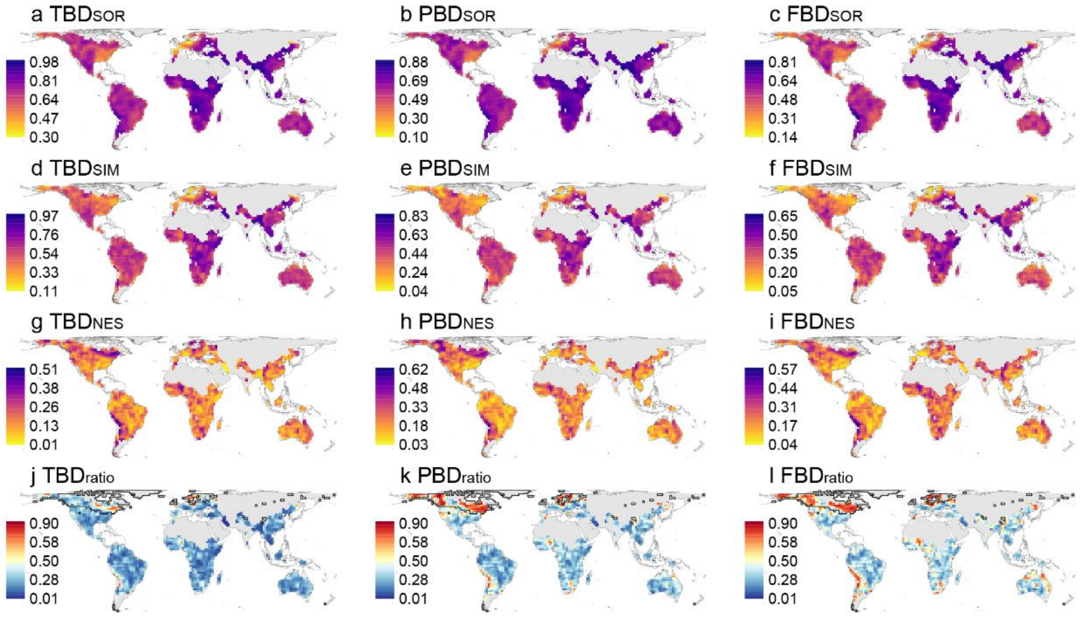


**Supplementary Fig. 5 | Geographic patterns of beta-diversity calculated with alpha-hull ranges using the alpha parameter of 2**. **a-c**, Total beta-diversity for the taxonomic ($TBD_{SOR}$), phylogenetic ($PBD_{SOR}$), and functional ($FBD_{SOR}$) dimensions; **d-f**, the component attributable to spatial turnover ($TBD_{SIM}$; $PBD_{SIM}$, $FBD_{SIM}$); **g-i**, the component attributable to nestedness ($TBD_{NES}$; $PBD_{NES}$, $FBD_{NES}$); **j-l**, the proportion of total beta-diversity contributed by nestedness component ($TBD_{ratio}$; $PBD_{ratio}$, $FBD_{ratio}$). Black lines in j-l showed grid-cells with over half of its area covered by ice during the Last Glacial Maximum, calculated with updated Quaternary glaciation coverage maps. Grid-cells with fewer than five species or five adjacent cells are shown in gray.

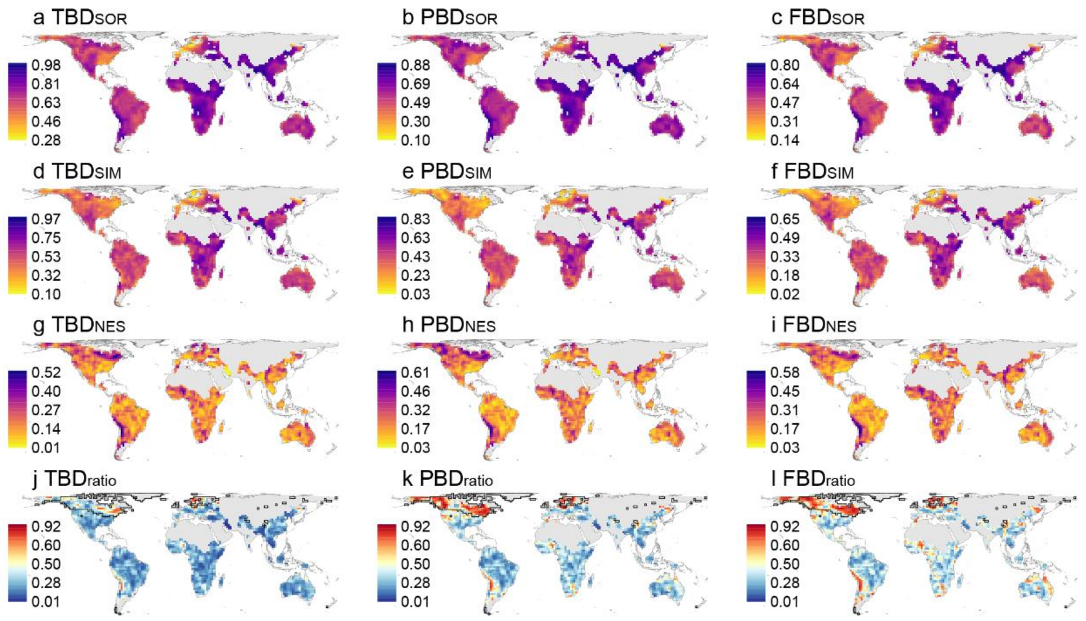


**Supplementary Fig. 6 | Geographic patterns of beta-diversity calculated with alpha-hull ranges using the alpha parameter of 4. a-c**, Total beta-diversity for the taxonomic ($TBD_{SOR}$), phylogenetic ($PBD_{SOR}$), and functional ($FBD_{SOR}$) dimensions; **d-f**, the component attributable to spatial turnover ($TBD_{SIM}$; $PBD_{SIM}$, $FBD_{SIM}$); **g-i**, the component attributable to nestedness ($TBD_{NES}$; $PBD_{NES}$, $FBD_{NES}$); **j-l**, the proportion of total beta-diversity contributed by nestedness component ($TBD_{ratio}$; $PBD_{ratio}$, $FBD_{ratio}$). Black lines in j-l showed grid-cells with over half of its area covered by ice during the Last Glacial Maximum, calculated with updated Quaternary glaciation coverage maps. Grid-cells with fewer than five species or five adjacent cells are shown in gray.

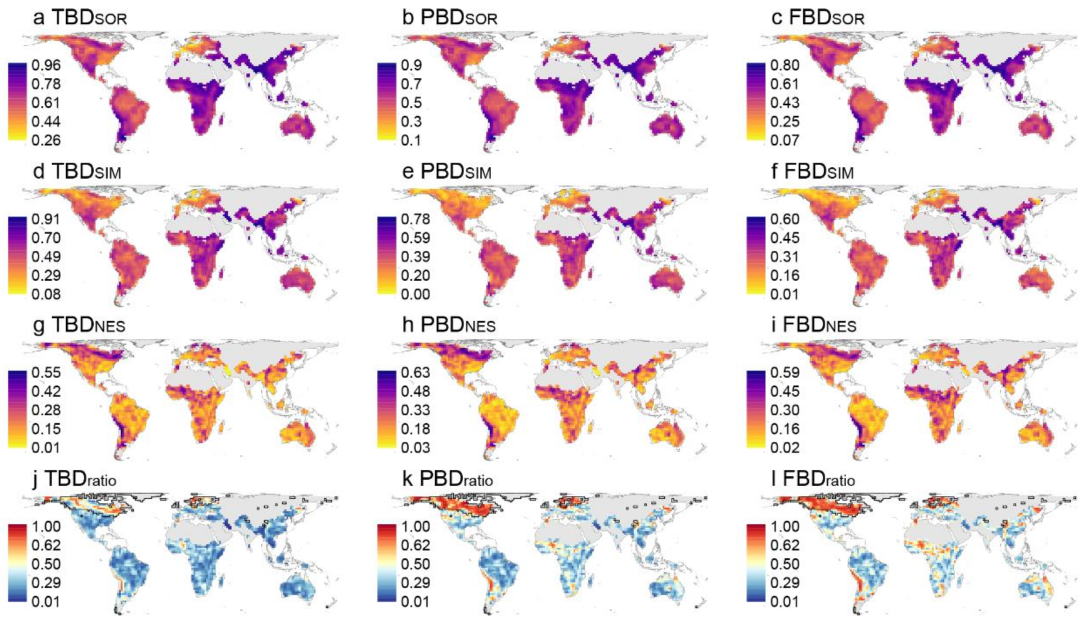


**Supplementary Fig. 7 | Geographic patterns of beta-diversity calculated with alpha-hull ranges using the alpha parameter of 10. a-c**, Total beta-diversity for the taxonomic ($TBD_{SOR}$), phylogenetic ($PBD_{SOR}$), and functional ($FBD_{SOR}$) dimensions; **d-f**, the component attributable to spatial turnover ($TBD_{SIM}$; $PBD_{SIM}$, $FBD_{SIM}$); **g-i**, the component attributable to nestedness ($TBD_{NES}$; $PBD_{NES}$, $FBD_{NES}$); **j-l**, the proportion of total beta-diversity contributed by nestedness component ($TBD_{ratio}$; $PBD_{ratio}$, $FBD_{ratio}$). Black lines in j-l showed grid-cells with over half of its area covered by ice during the Last Glacial Maximum, calculated with updated Quaternary glaciation coverage maps. Grid-cells with fewer than five species or five adjacent cells are shown in gray.

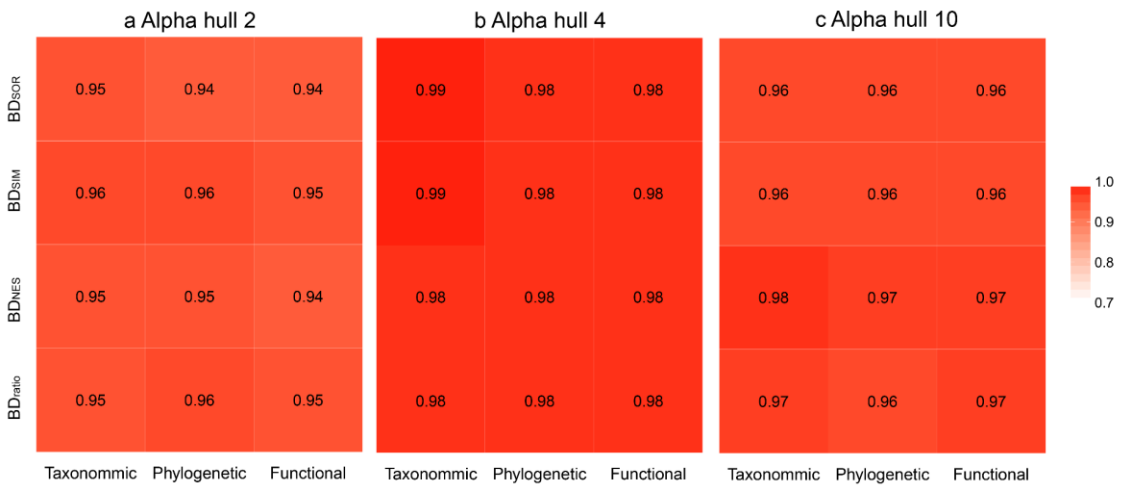


**Supplementary Fig. 8 | Pearson's correlations between beta-diversities calculated with alpha-hull ranges using different values of the alpha parameter.** The beta-diversity using the alpha parameter of 6 were compared to beta-diversities using the alpha parameter of 2 (**a**), 4 (**b**) and 10 (**c**). The comparisons were performed for taxonomic, phylogenetic and functional total beta-diversity ($BD_{SOR}$) and their respective components of turnover ($BD_{SIM}$) and nestedness ($BD_{NES}$) and the proportion of total beta-diversity contributed by nestedness ($BD_{ratio}$), respectively.

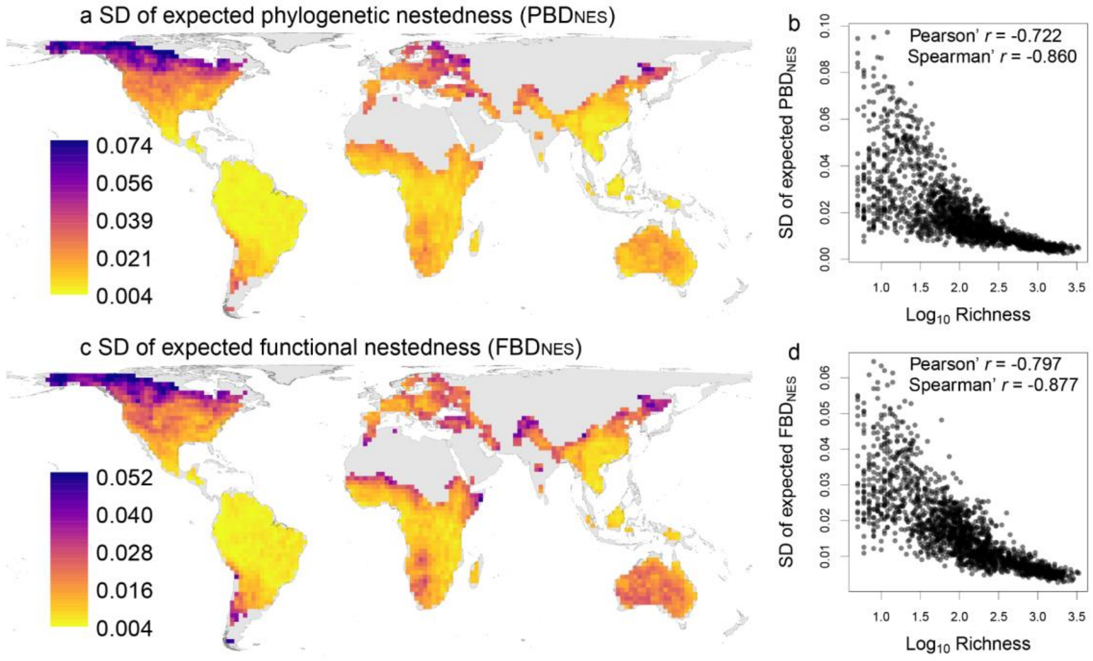


**Supplementary Fig. 9 | Geographic patterns of the standard deviation (SD) of randomly expected phylogenetic and functional nestedness (a, c) and relationships with species richness (b, d).** Both Pearson's and Spearman's correlation coefficients were provided in b and d. Note, because of strong relationships between the SD of expected beta-diversity and species richness, and striking spatial variations of species richness and the SD, the deviation of observed beta-diversity from the random expectation was not standardized by the SD of expected values. Grid-cells with fewer than five species or five adjacent cells are shown in gray in a and c.

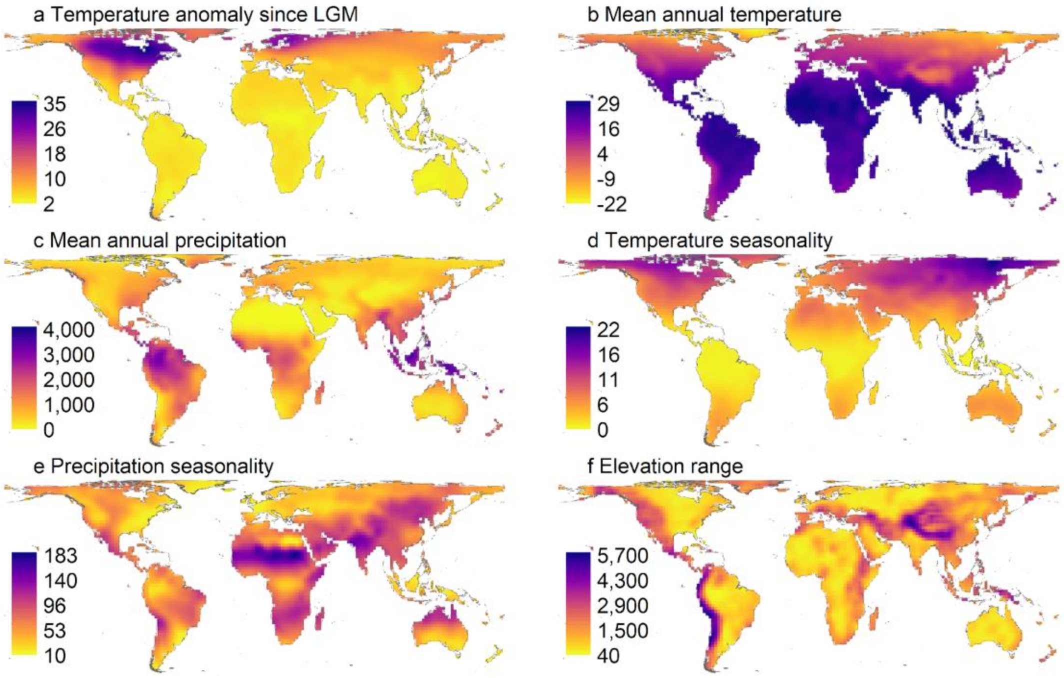


**Supplementary Fig. 10 | Geographic patterns of six environmental variables used in this study at the resolution of 200 × 200 km.**

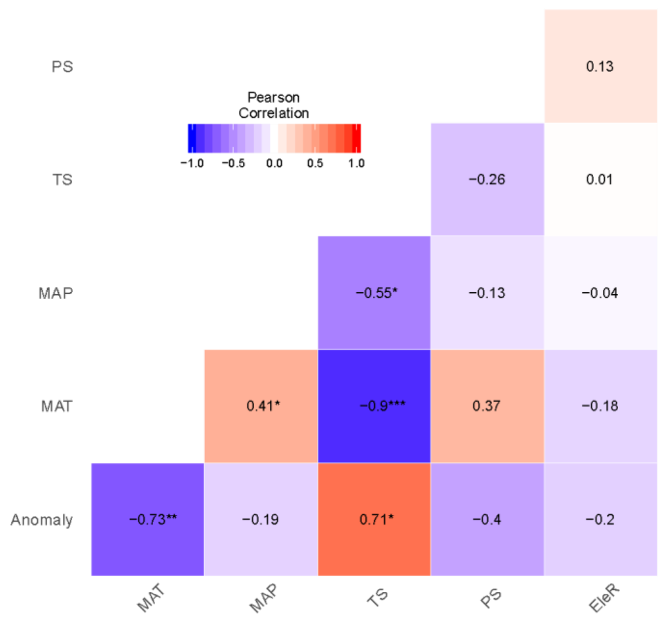


**Supplementary Fig. 11 | Pearson's correlations among six environmental variables.** Significance was tested using modified *t*-test to control for spatial autocorrelation. Anomaly, temperature anomaly since the Last Glacial Maximum; MAT, mean annual temperature; MAP, mean annual precipitation; TS, temperature seasonality; PS, precipitation seasonality; EleR, elevational range within grid-cells. *, $P < 0.05$; **, $P < 0.01$; ***, $P < 0.001$.

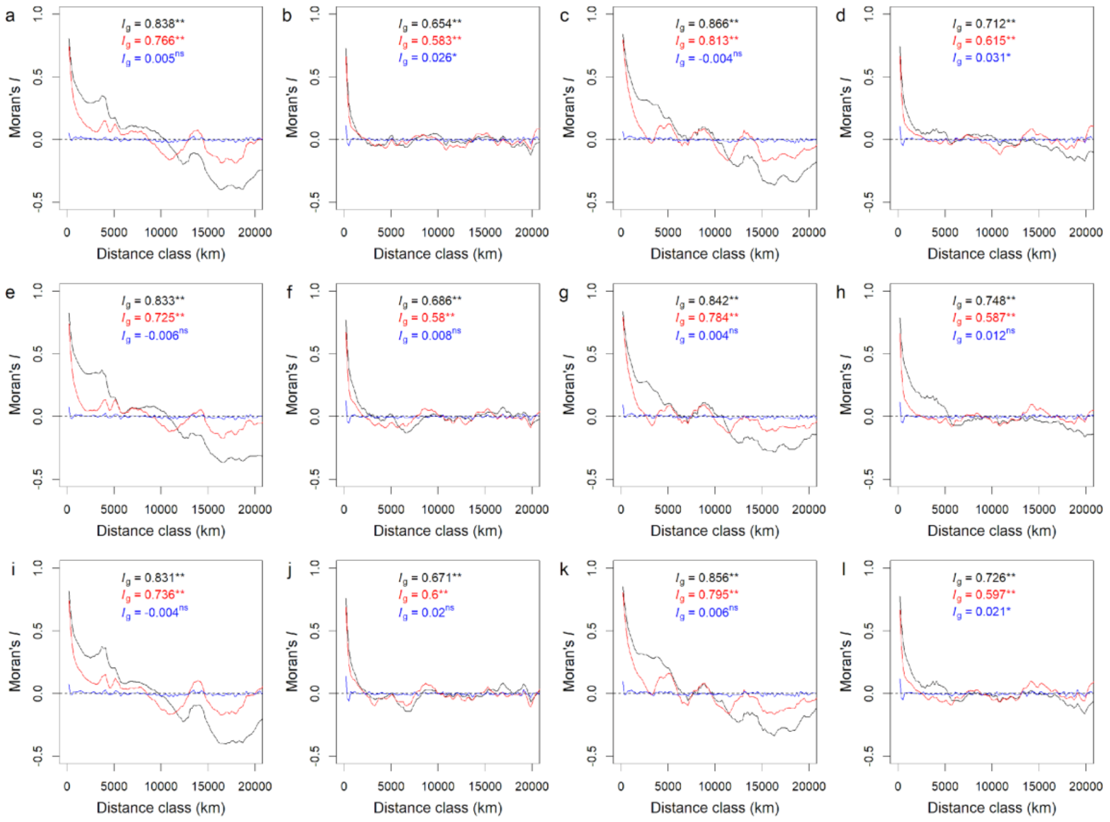


**Supplementary Fig. 12 | Spatial autocorrelation structures (Moran's *I*) of observed beta-diversity (black), residuals of full OLS models (red) and full SAR models (blue).** The results were shown for total beta-diversity (a, e, i) and the components attributable to spatial turnover (b, f, j) and nestedness (c, g, k) and the proportion of total beta-diversity contributed by nestedness component (d, h, l) for the taxonomic (a-d), phylogenetic (e-h) and functional (i-l) dimensions. The values in each panel show the Global Moran's *I* value ($I_g$), which was calculated with a neighbor distance of 300 km. *, $P < 0.05$; **, $P < 0.01$; ***, $P < 0.001$; ns, not significant.
